# Post-transcriptional regulatory mechanisms in human islets inferred from cell type-specific eQTLs detected by single-cell RNA-seq

**DOI:** 10.1101/2025.01.21.633530

**Authors:** Twan J.J. de Winter, Miha Sovrovic, Han Sun, James D. Johnson, Patrick E. MacDonald, Anna L. Gloyn, Françoise Carlotti, Eelco J.P. de Koning, Anna Alemany

**Affiliations:** Department of Anatomy and Embryology, Leiden University Medical Center, Leiden, The Netherlands; Department of Internal Medicine, Leiden University Medical Center, Leiden, The Netherlands; Department of Pediatrics, Division of Endocrinology, Stanford School of Medicine, Stanford, CA, USA; Department of Cellular and Physiological Sciences, Life Sciences Institute, University of British Columbia, Vancouver, BC, Canada; Vancouver Coastal Health Research Institute, Vancouver, BC, Canada; Department of Pharmacology, University of Alberta, Edmonton, AB, Canada; Alberta Diabetes Institute, University of Alberta, Edmonton, AB, Canada; Stanford Diabetes Research Center, Stanford School of Medicine, Stanford, CA, USA; Department of Genetics, Stanford School of Medicine, Stanford, CA, USA

**Keywords:** eQTL, post-transcriptional regulation, 3′ UTR, type 2 diabetes, beta cells, miRNA, scRNA-seq

## Abstract

Gene regulatory networks (GRNs) must be robust to maintain cellular identity across individuals yet flexible to accommodate population genetic variants. Comparing expression quantitative trait loci (eQTLs) in health and disease can therefore reveal hidden players in gene regulation. Here, we developed a computational pipeline to infer post-transcriptional regulatory events and their functional consequences from eQTL signals located in the 3′ untranslated region of genes using single-cell RNA sequencing. As a case study, we repurposed datasets from human islets of donors with and without type 2 diabetes (T2D). We identified eQTL landscapes differing by cell type and diabetes status, with cell-type specificity associated with gene differential expression and RNA-binding proteins. Integrating eQTLs with GWAS and miRNA hits linked variants in *G6PC2*, *QDPR*, and *RELL1* to insulin secretion, endoplasmic reticulum stress, and oxidative phosphorylation. Our pipeline provides a framework to unravel post-transcriptional regulatory mechanisms in health and disease at cell type resolution.

## Introduction

Modulation of mRNA stability and degradation through the 3′ untranslated region (3′ UTR) is a common post-transcriptional gene regulation strategy to control mRNA levels in cells^1–3^. The 3′ UTR is a non-coding region of mRNA containing regulatory motifs that can be bound by RNA-binding proteins (RBPs) and non-coding RNAs, such as microRNAs (miRNAs). Modification of these regulatory motifs can alter binding events and, subsequently, the half-life of mRNAs^4^. Therefore, genetic variation in the 3′ UTR can influence gene expression levels^5^, thereby becoming cis-expression quantitative trait locus (eQTL).

Single-cell RNA sequencing (scRNA-seq) is a powerful and widely used technology for assessing cell-type-specific gene expression. However, traditional scRNA-seq studies often overlook genetically driven differences in gene expression due to a lack of genotype data. Here, we hypothesized that post-transcriptional regulatory mechanisms can be inferred from scRNA-seq data by investigating cell-type specific eQTLs located in the 3′ UTR. As a case study, we focused on pancreatic islet biology in the context of both health and disease, specifically type 2 diabetes (T2D). T2D, whose risk is linked to a combination of environmental, lifestyle^6,7^ and genetic factors^8^, is a common endocrine condition characterized by chronically elevated blood glucose concentrations caused by insufficient insulin secretion from pancreatic beta cells usually on a background of insulin resistance^7^. Its progression alters gene regulatory networks within the islets^9,10^, which has been extensively studied using scRNA-seq^11^.

To test our hypothesis, we repurposed human islet scRNA-seq datasets to extract 3′ UTR genetic variants and cell type-specific eQTLs utilizing the fact that standard protocols provide high 3′ UTR coverage^12^. By combining the landscape of 3′ UTR-specific eQTLs with GWAS studies, cell type specific RBP expression patterns or islet miRNA expression, we inferred post-transcriptional gene regulatory events in beta and alpha cells derived from donors with T2D and without diabetes (ND). Our results allowed us to assess functional allele-specific consequences on insulin secretion phenotypes and gene expression modules related to endoplasmic reticulum (ER) stress and mitochondrial metabolism. Thus, our pipeline leverages the combination of scRNA-seq and genetic variation to unravel the post-transcriptional regulatory roles of miRNAs and RBPs in health and disease.

## Results

### Detection of known and novel cell type-specific 3′ UTR eQTLs in human islets

We integrated 12 published Smart-seq and Smart-seq2 scRNA-seq datasets^13–22^ from human pancreatic islets that comprised a total of 90 donors without diabetes (ND) and 35 donors with type 2 diabetes (T2D) (Figure 1A, Table S1, Methods). We selected datasets generated using full-length RNA sequencing protocols rather than the commonly used 3′ capturing techniques (e.g., 10x Genomics Chromium) to ensure adequate read depth for genotyping^12^. The resulting dataset (Figure 1B and Figure S1A-E) contained canonical pancreatic cell types such as endocrine (i.e., alpha, beta, delta, PP/epsilon), exocrine (i.e., acinar, ductal), pancreatic stellate, endothelial, and immune cells. We also identified two additional clusters, cycling alpha (*GCG*^+^, *MKI67*^+^, *CDK1*^+^) and alpha/beta cells (*INS^+^, GCG^+^*). These were not reported in the original publications, where the corresponding cells were mostly annotated as alpha cells or left unassigned (Figure S1F), nevertheless became detectable after dataset integration and are consistent with clusters observed in larger datasets that use the 10x Genomics Chromium technology^23^.

**Figure 1.**
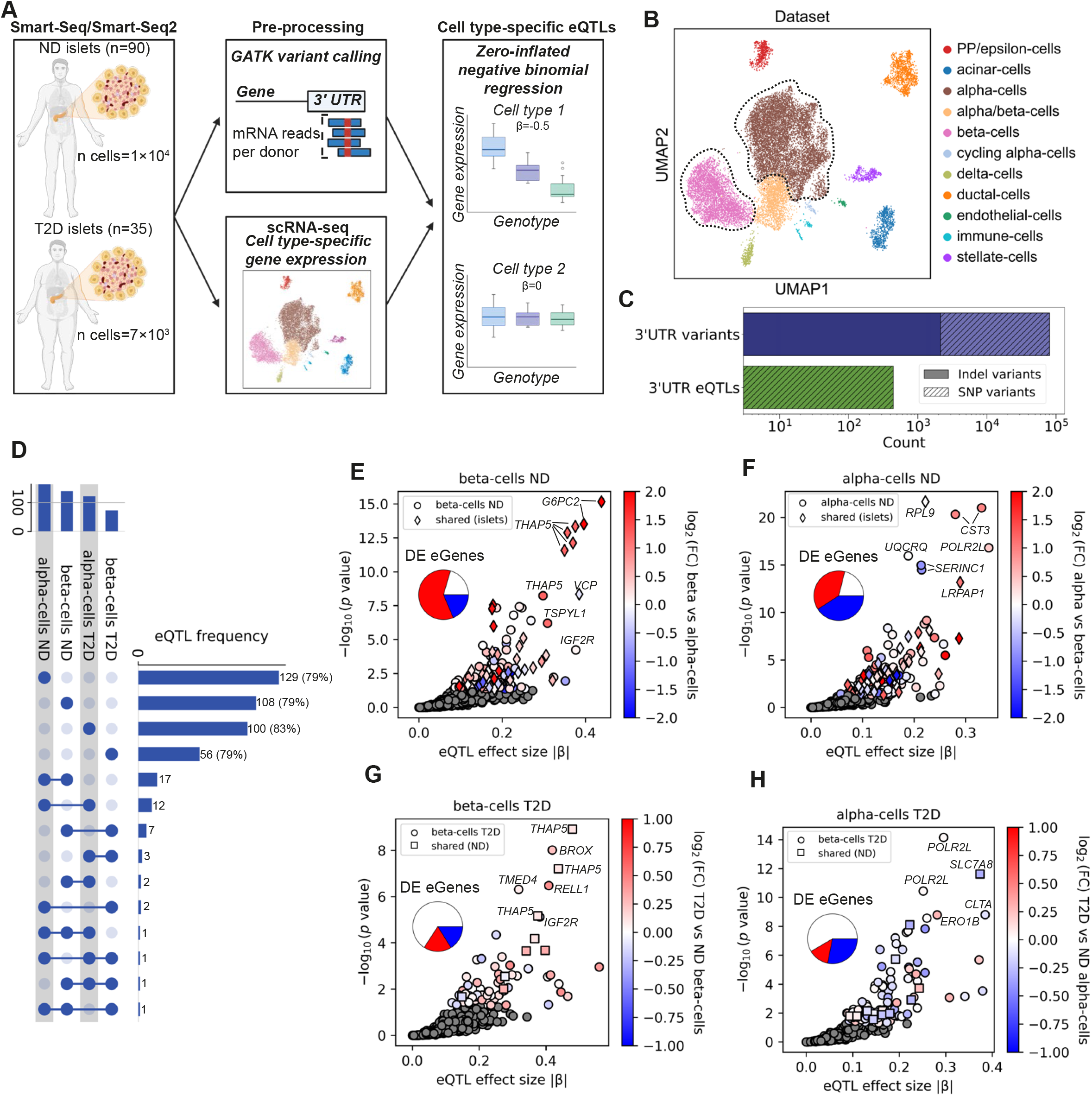
Detection of 3′ UTR eQTLs in human endocrine cell types. **A**, Schematic overview of the 3′ UTR eQTL detection pipeline from scRNA-seq data including 90 ND and 35 T2D donors. Created with BioRender.com. **B**, Uniform manifold approximation and projection (UMAP) of all cells, colored by cell type. Alpha and beta cell clusters used for eQTL detection are outlined with dashed lines. **C**, Bar graph showing the amount of detected 3′ UTR variants (minor allele frequency ≤ 0.07) and eQTLs in human islets. Darker and lighter colours with diagonal hatching indicate indel or SNP variants, respectively. **D**, Upset plot of eQTL frequency for alpha and beta cells from ND and T2D donors. Horizontal bars are annotated with eQTL frequency. **E-H**, Volcano plots of eQTL associations in beta and alpha cells from ND and T2D donors. Plots show absolute eQTL effect size (|β|) versus −log_10_(BH-adjusted p-value) from ZINB regression for beta cells (E,G) and alpha cells (F,H) in ND donors (E,F) and T2D donors (G,H). Significant eQTLs (BH-adjusted p < 0.05) are colored by log_2_ fold change (log_2_FC) between alpha and beta cells in ND donors for E,F, and between ND and T2D donors within the same cell type for G,H. Gray points indicate non-significant variant-gene pairs. Pie charts summarize eGenes as non-significant (white), upregulated (red), or downregulated (blue) in the corresponding comparison. Diamonds indicate eQTLs also detected in whole islets from the TIGER database; squares indicate T2D eQTLs also detected in the same cell type in ND donors. Top variant-gene pairs are annotated with gene symbols.

Next, we used the sequencing reads to genotype the common genetic variants (minor allele frequency ≥ 0.07; Methods) in each donor. We exclusively focused on genetic variants in the 3′ UTR because of this region’s important role in post-transcriptional regulation of gene expression^2^. Our integrated dataset contains 80,486 3′ UTR variants (Figure 1C), most being single nucleotide polymorphisms (SNPs) (97%) and a small amount (3%) consisting of insertions or deletions (indels). To identify which variants might regulate the expression of their host gene in a cell type-specific manner, thus behaving as cis-eQTLs (hereafter called eQTLs), we correlated the genotype with total gene expression level per cell. For each cell type we performed a zero-inflated negative binomial regression^24^, accounting for donor and cell-level fixed variables (Figure 1A, Methods). We stratified the analysis for the T2D status of the donors, and we only kept eQTLs exhibiting a monotonic allele-dosage effect on sufficiently expressed genes. Because the number of eQTLs detected in a certain cell type generally correlates with the number of cells interrogated^25,26^, we limited the analysis to the two most abundant cell types present in our integrated dataset, beta and alpha cells (n = 4.1×10^3^ and n = 8.7×10^3^ respectively, Figure 1B and Figure S1G).

A total of 440 variants act as an eQTL on 253 genes (hereafter called eGenes) in one or both cell types (Figure 1C,D and Table S2). We assessed whether the observed 3′ UTR eQTL effects could reflect linkage disequilibrium (LD) with active islet promoter or enhancer variants outside the 3′ UTR (Methods). The largest potential contribution was observed in ND alpha cells, where ∼14% of 3′ UTR eQTLs were in high LD with enhancer variants (*r*^2^ > 0.8, Figure S1H). Across all cell groups, high-LD promoter variants were less frequent than enhancer variants or absent. Our analysis revealed that eQTL effects were highly cell-type and T2D donor-status specific (Figure 1D). In ND beta cells, this can be attributed to the fact that ∼60% of eGenes are significantly upregulated in this cell type when compared to ND alpha cells (Figure 1E). However, the same reasoning does not hold for ND alpha cells (Figure 1F). Further exploration showed that only a few eQTLs were shared between ND and T2D cells (indicated as squares in Figures 1G,H), with T2D-specific eQTLs mostly present in eGenes not significantly differentially expressed between the disease status (Figure 1G,H).

We next checked whether the detected eQTLs had been previously characterized as whole islet eQTLs using the TIGER database (indicated as diamonds in Figures 1E, F), which contains whole islet eQTL information derived from 514 donors^27^. Overall overlap with whole islet signals was modestly higher in beta cells (36%) than in alpha cells (31%), although this difference was not significant (Fisher’s exact test p = 0.46, odds ratio [OR] = 0.82). Furthermore, we assessed how many of the eQTLs are also observed in bulk tissues from the Genotype-Tissue Expression (GTEx) database^28^ (Figure S1I), which includes whole pancreas samples. As expected, whole islets (TIGER database) are among the tissues with the highest overlap (34.3%). Consistent with the predominance of the exocrine compartment, only 16.6% of eQTLs are shared with whole pancreas eQTLs, and 13 non-pancreatic tissues show a larger overlap. Thus, our pipeline captures ∼66% of islet eQTLs previously missed by bulk islet transcriptomics studies.

Collectively, these findings demonstrate that 3′ UTR eQTLs display substantial cell type and disease-dependent differences. This pattern could be explained by differential expression (DE) of the associated eGenes in ND beta cells, but not in the other cell groups, highlighting a putative role of differential post-transcriptional regulatory landscapes in islets.

### Colocalization with islet-relevant GWAS variants reveals functional phenotypes of eQTLs

To date, over 600 T2D risk loci have been identified by genome-wide association studies (GWAS)^29^, some of which may reflect alpha or beta cell-specific eQTL effects. To investigate this possibility, we performed colocalization analysis between our list of cell-type-specific eQTLs and GWAS signals for T2D risk^29^ and glycemic traits, including glucose, insulin, and glycated hemoglobin levels (HbA1c; a measure of long-term blood glucose levels)^30^. The analysis aims to determine whether eQTL signals within the 3′ UTR of each eGene colocalize with GWAS signals, indicating that both associations are likely driven by a shared lead causal variant. Noteworthily, this analysis only gives significant results if the 3′ UTR contains enough variants for statistical analysis.

We found 9 eQTLs that colocalized with GWAS signals, none of them in LD with promoter or active islet enhancers. Among the eQTLs associated with glycemic traits (Figure 2A), we found two variants of *G6PC2* that are in moderate LD (*r*^2^ = 0.73; rs570876, chr2:168908867; rs567243, chr2:168909227), which are also the strongest ND beta cell eQTL signals in our dataset (Figure 1E). The *G6PC2* locus contains validated coding and non-coding variants that influence *G6PC2* expression, mRNA splicing, and protein function, and are associated with fasting blood glucose levels^31,32^. The colocalization analysis prioritizes rs567243 as the causal variant for fasting glucose^30^ (p = 2.6×10^-36^; Figure 2A,B) and rs570876 for HbA1c^30^ and T2D risk^29^ (p = 7.8×10^-6^ and p = 3.5×10^-5^; respectively, Figure 2A). G6PC2 is a negative regulator of glucose-stimulated insulin secretion (GSIS) that counteracts the activity of the main beta cell glucose sensor, GCK (Figure 2C)^33^. We therefore stratified GSIS phenotypes from the HumanIslets.com project^34,35^ by rs567243 genotype to assess whether the eQTL signal (Figure 2D) translates into altered insulin secretion. Although no association was found with the stimulation index in static GSIS after covariate adjustment (79 ND donors; Figure 2E), we observed significant associations in dynamic GSIS experiments (76 ND donors; Figure 2F, Figure S2A-D). These experiments included three sequential stimuli and captured the biphasic nature of insulin secretion. Consistent with static GSIS, rs567243 did not affect the secretory dynamics of the initial 15 mM glucose stimulus (Figure S2A,2D).

**Figure 2.**
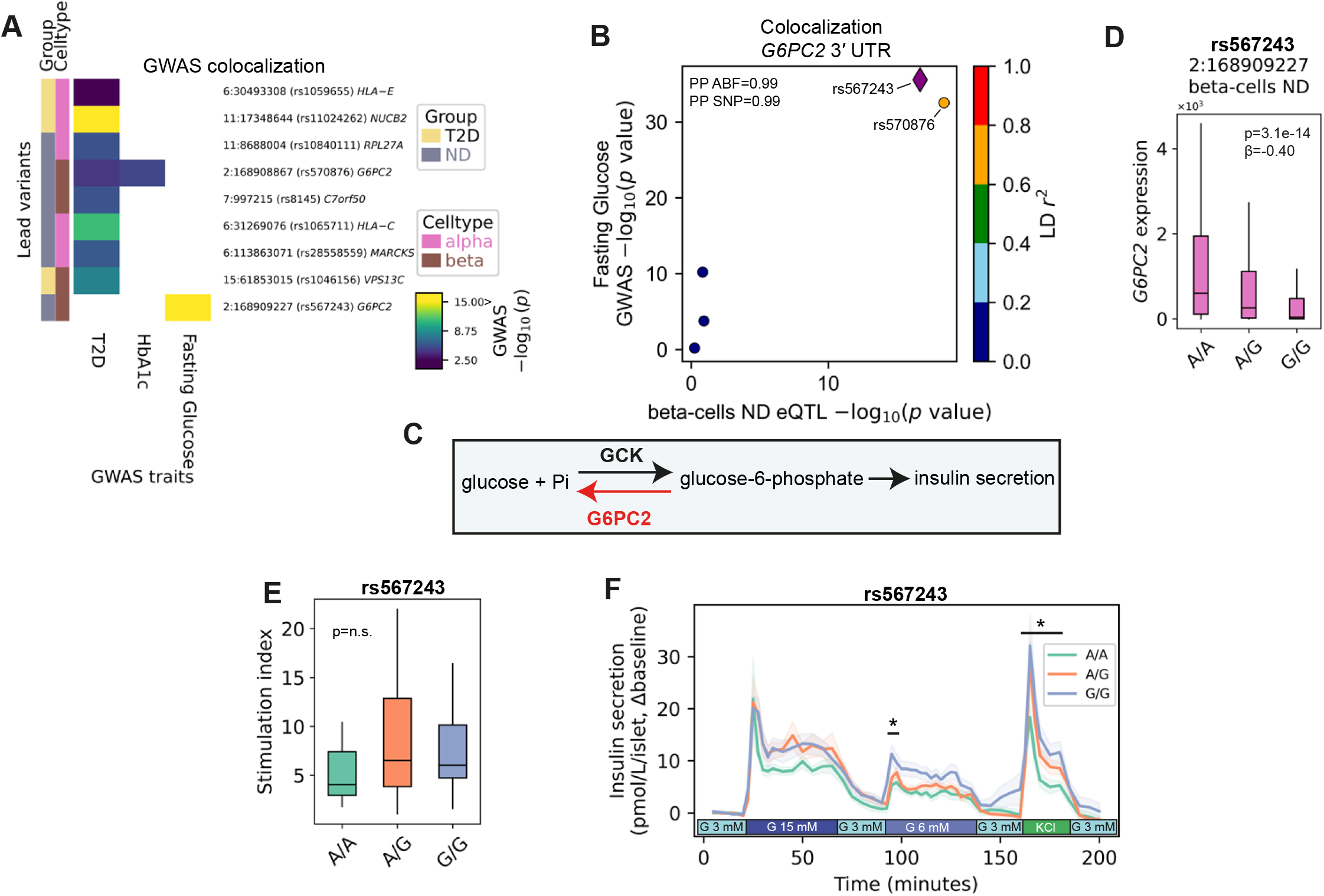
Colocalization of alpha and beta cells eQTLs with T2D risk and glycemic traits. **A**, Heatmap showing GWAS-eQTL colocalization for T2D and glycemic traits. The color indicates the −log_10_(p-value) of the GWAS association. Lead variants are annotated with cell type and diabetes status. **B**, LocusCompare-style plot showing the *G6PC2* eQTL-GWAS colocalization signal for fasting glucose. Shared 3′ UTR variants at the *G6PC2* locus are plotted by ND beta cell eQTL and GWAS −log_10_(p-value). The lead variant is highlighted by a purple diamond, and other variants are colored by LD (*r*^2^) with the lead variant. The plot also reports the coloc H4 posterior probability (PP ABF) and the lead SNP posterior probability (PP SNP). **C**, Schematic overview of the opposing enzymatic activities of GCK and G6PC2 to initiate glucose-stimulated insulin secretion (GSIS) in beta cells. **D**, Genotype-stratified boxplot showing *G6PC2* DEseq-normalized expression by rs567243 genotype in beta cells from ND donors. The plot is annotated with the BH-adjusted p-value from ZINB regression and eQTL effect size (β). Center line: median, box: interquartile range, vertical lines: spread of the data. **E**, Genotype-stratified boxplot showing the stimulation index (i.e., the ratio of insulin secreted in high-glucose to that in low-glucose) of ND donor islets assessed using static GSIS (low glucose, 1 mM; high glucose, 16.7 mM) by genotype of *G6PC2* variant rs567243. The plot is annotated with the p-value from linear regression adjusted for donor and islet isolation-level covariates. Center line: median, box: interquartile range, vertical lines: spread of the data. **F**, Genotype-stratified plot showing the average insulin secretion measurements of ND donor islets assessed using dynamic GSIS (basal glucose, 3mM; high glucose, 15 mM; moderate glucose, 6 mM; 30 mM KCl) by genotype of *G6PC2* variant rs567243. The shaded lines represent the SEM. Significant peaks and/or AUCs are marked with an asterisk.

Nevertheless, A/A carriers, associated with higher *G6PC2* expression (Figure 2D), exhibit a lower second-phase response (Figure 2F). Notably, increasing A-allele dosage was associated with a blunted first-phase peak during the subsequent 6 mM glucose stimulus (p = 4.8×10^-2^; Figure 2F, Figure S2B) and decreased KCl-induced, glucose-independent insulin secretion (Figure 2F, Figure S2C,F), consistent with differences in secretory capacity and/or total insulin content. In sum, our results suggest that the rs567243 A-allele is associated with increased *G6PC2* expression and reduced donor islet insulin output, which may contribute to elevated blood glucose levels.

All other identified lead variants were associated with T2D risk and included eQTLs from both ND and T2D alpha and beta cells. Collectively, colocalization analysis allowed to infer potential functional phenotypes of eQTLs located in the genes’ 3′ UTR using information from GWAS signals. However, the molecular mechanism connecting the eQTL to the phenotype is still missing. For this reason, we next aimed to investigate whether eQTLs where enriched in RBP and miRNA binding sites.

### Islet 3′ UTR eQTLs are enriched in RNA-binding protein motifs

To investigate whether RBP expression profiles can contribute to the observed eQTL cell-type specificity, we performed DE analysis between alpha and beta cells restricted to genes encoding RBPs (Table S3)^36,37^. A substantial fraction of RBPs showed clear DE signatures between ND alpha and ND beta (63% of RBPs, Figure 3A), indicating distinct post-transcriptional machinery between the two cell types. In line with previous literature, we identified *RBFOX3* and *ELAVL4* highly upregulated RBPs in beta cells and *ZFP36* highly expressed in alpha cells^38–40^.

**Figure 3.**
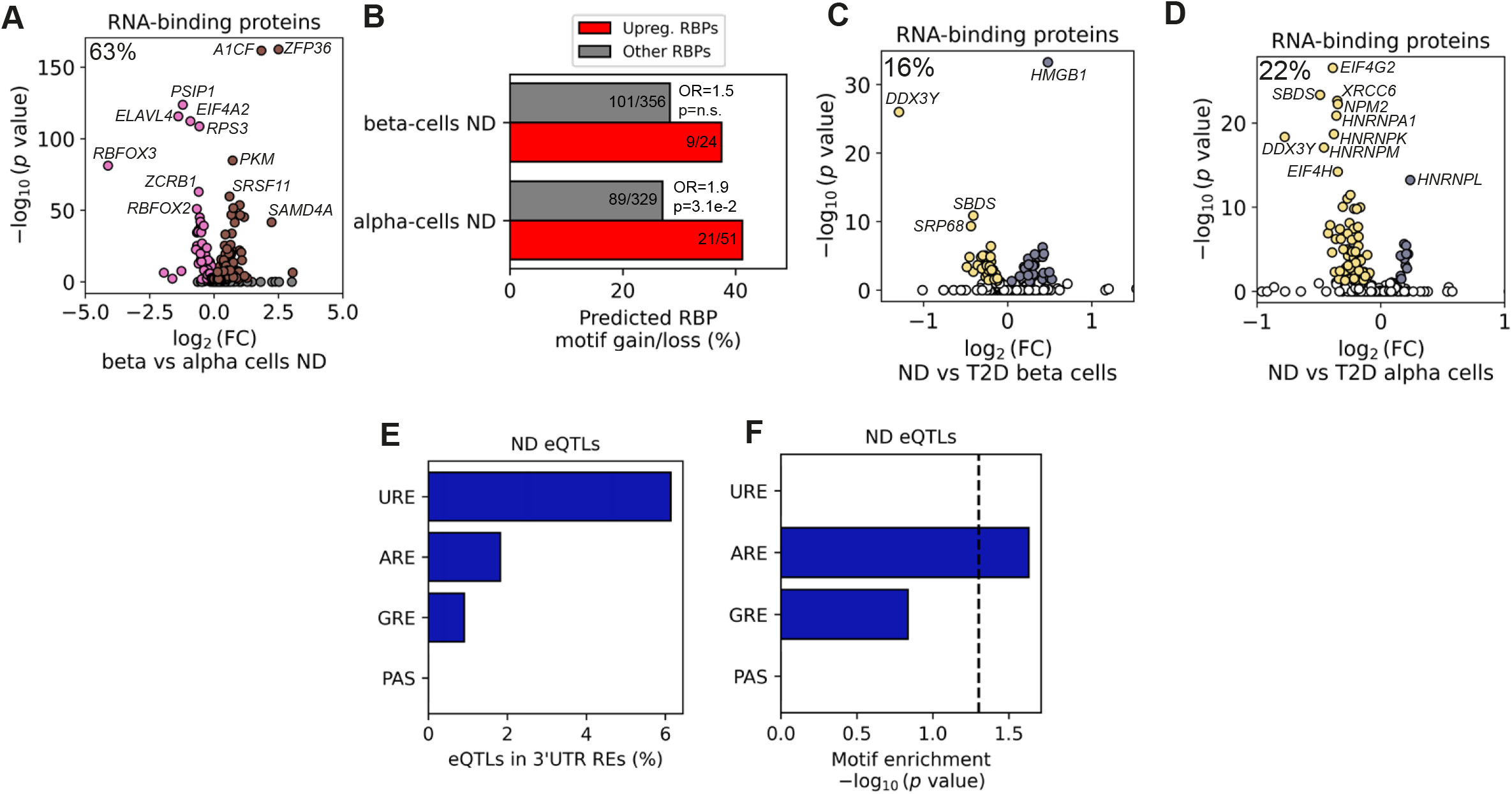
Cell type specific eQTLs are enriched for differentially expressed RBPs and stabilizing 3′ UTR elements. **A**, Volcano plot of differential expression of RNA-binding proteins (RBPs) genes between beta and alpha cells in ND donors, showing log_2_(FC) versus −log_10_(BH-adjusted p-value) from a Wilcoxon rank-sum test. Pink and brown points indicate RBPs upregulated (p < 0.05) in beta and alpha cells, respectively. Gray points indicate non-significant RBPs. The plot is annotated with the percentage of RBPs that are differentially expressed. Top RBPs are annotated with gene symbols. **B**, Horizontal bar graph showing the percentage of upregulated (Upreg.) and the remaining RBPs in the dataset (Other) where ND eQTL variants are predicted to alter the binding motif (motif gain/loss), per cell type. The plot is annotated with the p-value and odds ratio (OR) from a Fisher’s exact test. **C**,**D**, Volcano plots of differential expression of RBP genes of beta (C) and alpha cells (D) between ND and T2D donors within the same cell type, showing log_2_(FC) versus −log_10_(BH-adjusted p-value) from a Wilcoxon rank-sum test. Yellow and gray points indicate RBPs upregulated (p < 0.05) in ND and T2D cells, respectively. White points indicate non-significant RBPs. The plot is annotated with the percentage of RBPs that are differentially expressed. Top RBPs are annotated with gene symbols. **E**,**F**, Horizontal bar graphs showing the percentage (E) and motif enrichment (−log_10_ p-value) (F) of ND eQTLs that are located within 3′ UTR regulatory elements (RE): U-rich element (URE), AU-rich element (ARE), G-rich element (GRE), and polyadenylation signal (PAS). The vertical dashed line marks significance (p < 0.05).

We reasoned that RBPs’ DE can shape the cell-type specific eQTL landscape if the identified 3′ UTR eQTL variants alter RBP-binding motifs. Therefore, we set up to investigate whether this is the case (Methods). Out of the 380 RBP detected in our scRNAseq dataset, we identified 120 for which eQTL variants were predicted to affect binding affinity. In alpha cells, RBPs with predicted altered binding were enriched among those upregulated in alpha cells (Fisher’s exact test p = 3.1×10^-2^, OR = 1.9; Figure 3B). RBP expression for each cell type between T2D-status revealed only modest changes (17% in beta cells and 22% in alpha cells; Figure 3C,D). Altogether, this indicates that these two cell types harbour distinct post-transcriptional machinery, which might explain the eQTL cell type specificity.

However, these results argue against large-scale changes in RBP abundance as the primary driver of this disease-dependent eQTL remodeling.

To further explore the role of RBPs in eQTL regulation, we assessed whether eQTL variants influence four common 3′ UTR regulatory elements (RE) bound by RBPs, namely AU-rich (ARE), GU-rich (GRE), U-rich (URE), and canonical polyadenylation signal (PAS) motifs (Figure S3A)^41^. AREs, present in 56% of mRNAs, are well-established mediators of mRNA stability^42,43^. URE and GRE motifs serve broader regulatory roles and are enriched near 3′ UTR cleavage sites, where they are essential for 3′ UTR end formation^44,45^. We found only a small subset of eQTLs altering motifs of annotated 3′ UTR REs, with most affecting UREs (Figure 3E), consistent with their high abundance^46^. None of the identified eQTLs overlapped PAS motifs, essential for 3′ UTR cleavage and polyadenylation. Strand-aware motif enrichment analysis (Methods) revealed that only ARE motifs were significantly overrepresented within and immediately surrounding eQTL sites (Figure 3F, S3B). Notably, for URE and PAS motifs, we observed sharp enrichment peaks of about 25 nucleotides away from eQTL positions (Figure S3B), suggesting that eQTLs proximal to URE and PAS motifs may influence supporting regulatory motifs, such as the cleavage factor Im (CFIm) binding motif UGUA^47^. Collectively, our results suggest that eQTLs are enriched in mRNA stabilizing motifs.

### 3′ UTR eQTLs predict allele-dependent changes in islet miRNA binding

Next, we investigated whether 3′ UTR variants could give rise to eQTLs by altering miRNA binding sites. To do this, we first measured the miRNA expression profiles of whole islets (islet tissue purity >85%) from ND and T2D donors (n = 6 for both) using small RNA-sequencing (Figure 4A, Table S4, and Methods). After filtering, we obtained the expression profile of 543 miRNAs (Figure S4A,B), the most highly expressed being members of the let-7 miRNA family and hsa-miR-375-3p (Figure S4B).

**Figure 4.**
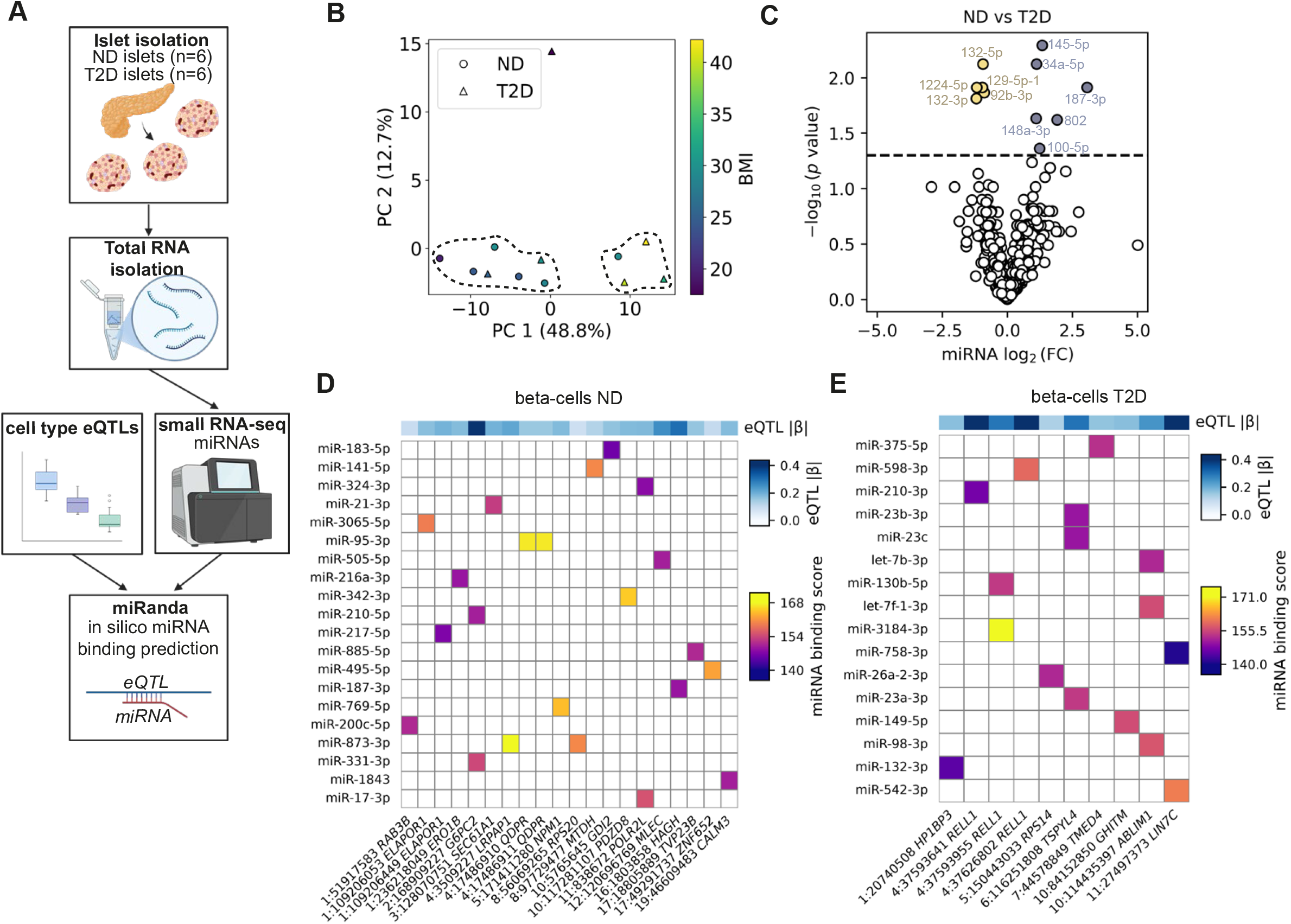
A subset of eQTLs is predicted to alter miRNA binding sites. **A**, Schematic overview of the small RNA-sequencing pipeline containing 6 ND and 6 T2D donors for miRNA expression analysis and binding prediction. Created with BioRender.com. **B**, Prinicipal component analysis (PCA) of islet miRNA expression profiles. The scatter plot shows the first versus second principal component (PC). PC1 and PC2 explain 48.8% and 12.7% of total variance, respectively. Points are colored by the BMI of the donor, and the shape indicates the diabetes status. Main cluster separation is outlined with dashed lines. **C**, Volcano plot showing log_2_(FC) of islet miRNAs versus −log_10_(BH-adjusted p-value) obtained from differential miRNA expression analysis using the DEseq method. The horizontal dashed line marks significance (p < 0.05). Yellow and gray points indicate miRNAs upregulated in ND and T2D islets, respectively. Significant miRNAs are annotated. **D**,**E,** All predicted miRNA-eQTL pairs in beta cells in ND (D) and T2D donors (E) with disrupted binding sites. The top bar shows absolute eQTL effect sizes (|β|) of donors within the same cohort. The heatmap shows miRanda *in silico* binding scores.

Principal component analysis (PCA) showed moderate separation of donor islet miRNA expression by body mass index (BMI) and disease status (Figure 4B). Most ND samples clustered together with two T2D donors with moderate BMI values, whereas T2D donors with higher BMI/severe obesity were enriched in a separate cluster. One underweight T2D donor was separated from the remaining samples. Along these lines, only 11 miRNAs showed significant DE levels between ND and T2D donor’s islets (Figure 4C), 7 of which were also correlated with HbA1c (Figure S4D). Although BMI was significantly higher in T2D donors compared to ND (Student’s t-test p = 4.7×10^-12^), we did not find this to be a strong driving factor of the DE (Figure S4C).

The modest miRNA DE detected between ND and T2D samples let us to use the top 95% expressed islet miRNAs in the whole dataset (Figure S4B) to predict allele-specific binding gain and loss between miRNA seed regions and eQTL variants by computing miRNA binding scores using the miRanda algorithm^48^ (Figure 4A, Methods). For this analysis, we focused on the eQTLs detected in beta cells for both ND and T2D because this is the predominant cell type in islets. We infer that 36 miRNAs (none of the 11 DE between T2D and ND islets) can explain the behavior of 30 eQTLs in our dataset, with 20 and 16 miRNA-eQTL pairs for ND and T2D beta cells, respectively (Figure 4D,E; Figure S4E,F; Table S5). None of the miRNA-eQTL pairs were shared between ND and T2D. Thus, our results suggest that a subset of eQTLs in beta cells might originate through the creation or disruption of miRNA binding sites.

### Functional implications of allele-dependent miRNA regulation

The miRNA-eQTL pairs with the strongest *in silico* binding scores from ND beta cells correspond to two adjacent variants, rs1049581 and rs1049582, that alter the binding site of hsa-miR-95-3p (hereafter called miR-95; Figure 4D, S4B) in the 3′ UTR of *QDPR.* These two variants, in near-perfect LD (r^2^ > 0.99; chr4:17486910-17486911), are also collectively reported as rs386672163, which has previously been identified as an islet eQTL in the public TIGER database (p = 3.1×10^-4^)^27^. In line with the low expression levels of *QDPR* in beta cells from donors homozygous for the reference alleles (rs1049582: C; rs1049581: A; Figure 5A), the alternative alleles (rs1049582: T; rs1049581: G) disrupt miRNA binding (Figure 5B). We also scanned the full 3′ UTR sequence of *QDPR* and did not find any additional predicted miR-95 binding sites (Figure S5A).

**Figure 5.**
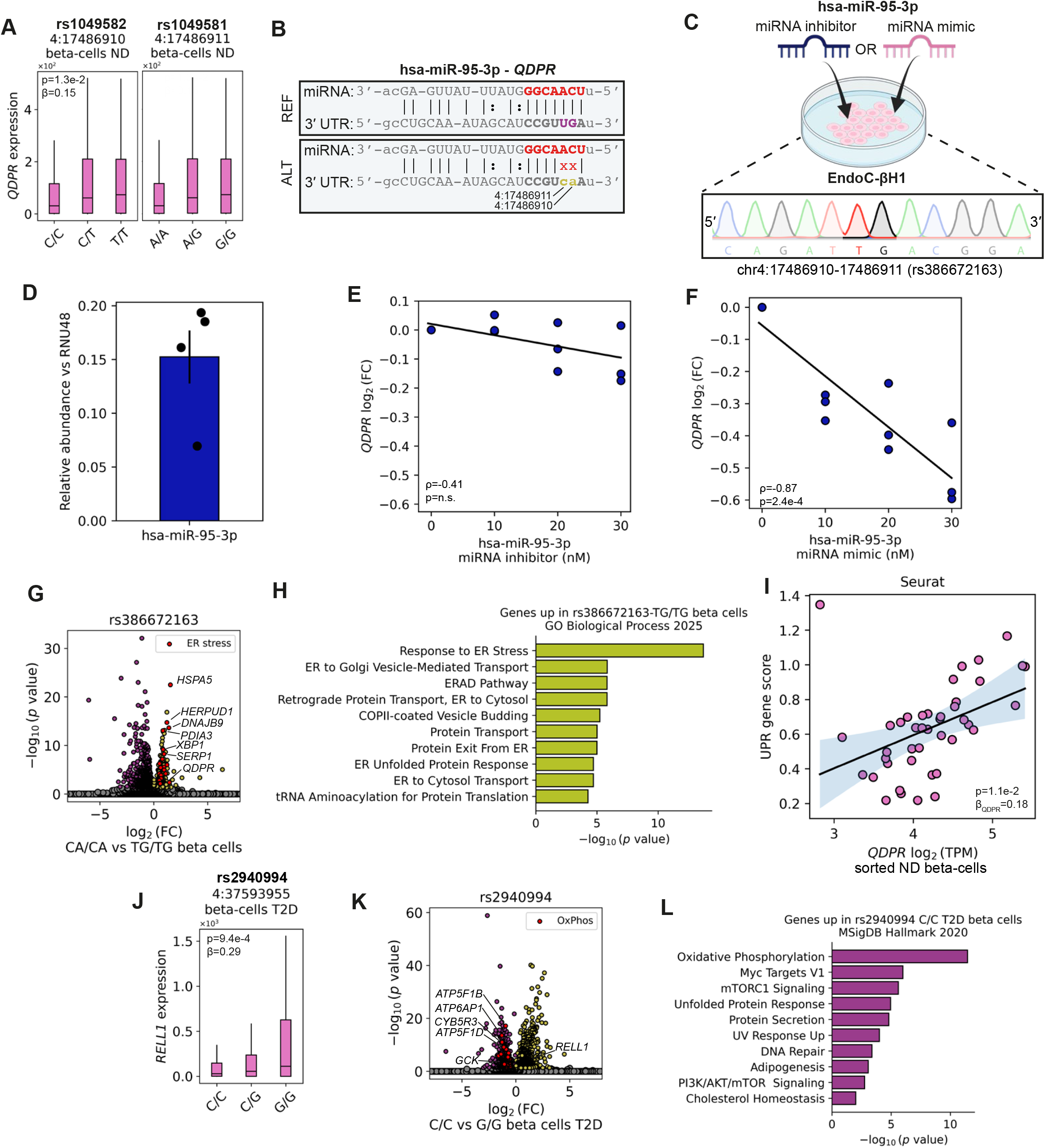
Validation of miRNA binding and assessment of functional consequences of altered miRNA binding. **A**, Genotype-stratified boxplots showing QDPR DEseq-normalized expression by rs1049581 and rs1049582 (collectively reported as rs386672163) genotype in beta cells from ND donors, annotated with the BH-adjusted p-value from ZINB regression and eQTL effect size (β). Center line: median, box: interquartile range, vertical lines: spread of the data. **B**, Nucleotide alignment of hsa-miR-95-3p to the 3′ UTR of QDPR shown in the orientation of transcription after Watson and Crick and Wobble base pairing. miRNA seed region shown in bold red, the reference allele (binding) in purple, and the alternative allele (disrupting) in gold, and annotated with the genomic position. **C**, Schematic overview of the miR-95-3p knockdown (inhibitor) and overexpression (mimic) experiment in EndoC-β1. Below is the Sanger sequencing chromatogram of the QDPR 3′ UTR showing the alternative genotypes of rs386672163 (chr4:17486910-17486911) displayed in genomic orientation and therefore as the reverse complement of panel B. **D**, Relative abundance of miR-95-3p in EndoC-β1 normalized to the housekeeping snoRNA RNU48. N = 4 biological replicates, mean ± SEM. **E**,**F**, Scatter plot showing the correlation between log2(GAPDH-corrected QDPR) abundance normalized to the non-transfection control measured by RT-qPCR versus increasing concentrations of miR-95-3p inhibitor (E) and mimic (F) transfected in EndoC-β1. N = 3 biological replicates, each point represents the average of three cell culture wells. Plots are annotated with the Rho (ρ) and p-value obtained from Spearman correlation analysis. The solid line indicates the result of a linear fit to the displayed data. **G**, Volcano plots of differential expression analysis of genes in ND beta cells between donors with homozygous genotypes of rs386672163, showing log2(FC) versus −log10(BH-adjusted p-value) from a Wilcoxon rank-sum test. Purple and gold points indicate genes upregulated (p < 0.05) in CA/CA (homozygous reference) and TG/TG (homozygous alternative) beta cells, respectively. Gray points indicate non-significant genes. Red points indicate genes part of the “Response to ER stress” module from panel H, with the top and QDPR annotated with gene symbols. **H**, Top 10 results from gene set enrichment analysis (GO biological process) using upregulated genes in beta cells with the alternative rs386672163 TG/TG genotype from panel G. **I**, Scatter plot showing the association between log2(normalized QDPR) expression versus UPR gene score in sorted beta cells. Gene score was calculated using Seurat, and the plot is annotated with the effect size (β) and p-value obtained from regression analysis adjusted for donor-level covariates. The solid line indicates the result of a linear fit to the displayed data. **J**, Genotype-stratified boxplot showing RELL1 DEseq-normalized expression by rs2940994 genotype in beta cells from T2D donors. The plot is annotated with the BH-adjusted p-value from ZINB regression and eQTL effect size (β). Center line: median, box: interquartile range, vertical lines: spread of the data. **K**, Volcano plots of differential expression analysis of genes in ND beta cells between donors with homozygous genotypes of rs2940994, showing log2(FC) versus −log10(BH-adjusted p-value) from a Wilcoxon rank-sum test. Purple and gold points indicate genes upregulated (p < 0.05) in C/C (homozygous reference) and G/G (homozygous alternative) beta cells, respectively. Gray points indicate non-significant genes. Red points indicate genes part of the “Oxidative Phosphorylation” (OxPhos) module from panel L, with the top, GCK, and RELL1 annotated with gene symbols. **L**, Top 10 results from gene set enrichment analysis (MsigDB Hallmark) using upregulated genes in beta cells with the reference rs2940994 C/C genotype from panel K.

We set up to validate this miRNA-mediated interaction using EndoC-βH1 cells, an immortalized human beta cell line homozygous for the disrupting alternative alleles (Figure 5C). Because this cell line has detectable levels of miR-95 (Figure 5D, S5B), we first performed knockdown of this miRNA. Consistent with our prediction, transfecting EndoC-βH1 with increasing concentrations of a synthetic miR-95 inhibitor did not affect *QDPR* expression (Figure 5E), suggesting limited miRNA-mediated repression. Conversely, while a non-targeting miRNA control did not affect *QDPR* expression (Figure S5C), miR-95 overexpression via a synthetic mimic significantly repressed *QDPR* in a dose-dependent manner (Spearman ρ = -0.87; p = 2.4×10^-4^; Figure 5F). Collectively, these findings proof the validity of our bioinformatic pipeline to infer miRNA-eQTL functional pairs and further confirm that endogenous miRNA-mediated repression of *QDPR* is minimal in EndoC-βH1 cells, consistent with impaired miRNA binding associated with the cell line’s genotype.

We next reasoned that we could use our integrated scRNA-seq dataset to investigate whether the *QDPR* eQTL has functional consequences in beta cells. *QDPR* encodes DHPR, a quinoid dihydropteridine reductase, a metabolic enzyme that recycles tetrahydrobiopterin (BH4) levels by converting quinoid-BH2^49^, and its role in beta cells is poorly understood. Although no GWAS traits have been reported for rs386672163, we found that genes upregulated in beta cells homozygous for the rs386672163 reference allele (CA/CA) compared to the alternative allele (TG/TG) (Figure 5G) were strongly enriched for ER stress and unfolded protein response (UPR) modules (Figure 5H, S5D, Table S6). To further investigate the possible link between *QDPR* and ER stress, we analyzed an independent publicly available bulk RNA-seq dataset of sorted human beta cells from 42 ND donors, none of whom were included in our eQTL dataset (Methods)^50^. *QDPR* expression was significantly correlated with UPR stress score determined using two independent gene-scoring methods (Methods, Seurat, p = 1.1×10^-2^; Singscore, p = 4.4×10^-4^; Figure 5I, S5E, Table S3). Importantly, *QDPR* expression was not associated with reduced beta cell identity (Figure S5F,G, Table S3). Thus, our results support a model in which rs386672163 acts through miR-95-dependent regulation, yielding allele-dependent differences in ER stress gene expression.

Lastly, we repeated genotype-aware DE analysis for the eQTL with the highest predicted miRNA binding score in T2D beta cells (Figure 4E). Our pipeline predicts that the alternative allele of the *RELL1* 3′ UTR variant rs2940994 (chr4:37593955) disrupts binding of hsa-miR-3184-3p (Figure 4C, 5J, S5H). Comparing homozygous rs2940994 genotypes (Figure 5K,L, Table S6) revealed that T2D beta cells carrying the reference C/C genotype significantly upregulated genes involved in oxidative phosphorylation, including core electron transport chain genes (e.g., *NDUFS2*, *UQCRC1*, *COX5A*, *ATP5F1A*) and citric acid cycle genes (e.g., *ACO2*, *OGDH*, *FH*). Notably, these cells also showed higher expression of *GCK*, the main beta cell glucose sensor involved in GSIS (Figure 2C). Conversely, this suggests that increased *RELL1* expression in alternative-allele carriers (Figure 5J) is associated with reduced expression of genes involved in glucose sensing and mitochondrial metabolism. This is consistent with *RELL1* encoding a membrane protein that positively regulates p38 MAPK, a signaling cascade implicated in oxidative stress, impaired GSIS, and apoptosis^51^. Together, these findings suggest that miRNA-based regulation of *RELL1* may contribute to a more pronounced beta-cell dysfunction in a genetically defined subset of T2D patients.

## Discussion

In this study, we show for the first time that cell type-specific post-transcriptional regulatory events can be detected from integrated scRNA-seq datasets containing many individuals. Accordingly, we provide a pipeline to accurately identify cell-type specific eQTLs and investigate whether these originate via the action of RBPs or miRNAs. To understand the functional consequences of post-transcription modification, we applied GWAS colocalization analysis and propose a genotype-aware DE analysis strategy for generating biological hypothesis regarding the role of specific post-transcriptional regulatory events.

Using pancreatic islet biology as a case study, we integrated previously published scRNAseq data from 90 donors without diabetes and 35 donors with T2D, from which we generated a novel eQTL database, consisting of 440 variants and uncovering distinct cell type and disease-specific eQTL behavior in endocrine cells. When comparing eQTLs in our study with those identified in whole islets by others, we only observed a ∼30% overlap^27^. This is consistent with reports from other tissues, including brain, blood, and skin, and aligns with the current consensus that bulk studies mask cell type- and context-dependent effects^25,52^. Our results suggest that this cell-type specificity is shaped by two complementary factors: the availability of the target transcript and the presence of post-transcriptional regulators in the relevant cell type. In beta cells, for example, most beta cell eGenes were absent or reduced in alpha cells. Conversely, in alpha cells, eQTLs predicted to alter RBP-binding motifs were enriched for RBPs with alpha-cell-specific expression. This finding is supported by previous work showing that the most differentially expressed RBPs play important roles in their respective cell types^38–40^, as well as by reports of distinct cell-type-specific mRNA splicing profiles^53^.

After identification and validation (when possible) of mRNA:miRNA pairs, performing genotype-aware DE analysis using the top miRNA-eQTL hits in ND and T2D beta cells showed possible functional implications for the predicted miRNA binding. For instance, we observed that ND donors carrying the alternative alleles of the *QDPR* rs386672163 variant have coordinated upregulation of an ER stress/UPR gene program, covering the canonical IRE1α-XBP1 axis (Table S6), with induction of downstream stress responders *DDIT3* (CHOP) and *PPP1R15A* (GADD34)^54^, which has not been reported before. In beta cells, nitric oxide synthase (NOS) has been shown to cause UPR stress^55^. Because QDPR provides cellular levels of BH4^56^, an essential cofactor of NOS^49^, we speculate that QDPR modulates beta cell nitric oxide levels, something that has been observed in other cell types^57^. In T2D donors, we showed that donors carrying the alternative allele of the *RELL1* variant rs2940994 have decreased expression of genes involved in oxidative phosphorylation and glucose sensing. RELL1 has been shown to activate the pro-apoptotic p38 MAPK signaling cascade^58^, a pathway implicated in sensing beta cell stress^51,59^. In mice, inhibition of p38 improves blood glucose levels, stability of the beta cell maturity marker MafA, and prevents apoptosis^51,60^. Thus, therapeutic modulation of RELL1-associated stress pathways may provide a targeted opportunity to reduce beta-cell stress in a genetically defined subset of T2D patients. Together, this shows that our pipeline can unravel biological processes related to post-transcriptional regulation.

Our study also has some limitations, such as the sample size for our eQTL analysis which is relatively modest compared to bulk studies. Additionally, for this study we genotyped islet donors from pseudobulk single-cell RNA-sequencing data, and this may lead to false variant calling, including among others, lowly expressed genes, genes with strong allele-specific gene expression, and mRNAs that are modified with endogenous RNA editors^61^. Moreover, the absence of genome-wide genotype information of most donors limits our ability to distinguish lead eQTL SNPs from correlated variants in LD. Finally, as isolated and cultured primary islets have been shown to induce transcriptional changes^35^, caution is warranted that some eQTL signals may arise ex vivo.

Regardless of these limitations, we showed that our computational approach gives insight into the post-transcriptional eQTL regulation landscape of human beta and alpha cells during normal physiology and T2D. Although we only applied our pipeline to single-cell datasets, we believe it can also be extended to bulk mRNA-seq datasets upon prior cell type purification. Thus, in the era of the human cell atlas, we envision that our workflow has the power to unveil post-transcriptional regulatory mechanisms in health and disease in many tissues and cell types with already available transcriptomics datasets.

## Methods

### Experimental model and subject details

#### Cell culture and maintenance

EndoC-βH1 cells (Human origin, female) were obtained from Human Cell Design. Cells were cultured in low glucose DMEM (Gibco, cat. no. 21885108), supplemented with human serum albumin (CSL Behring, cat. no. Rvg 105901, 2%), Nicotinamide (KFT-LUMC, cat. no. 97996807,10mM), Human Transferrin (Sigma-Aldrich, cat. no. T8158, 5.5g/mL), Selenite (Sigma-Aldrich, cat. no. S1382, 0.5g/mL),

Penicillin/Streptomycin (Gibco, cat. no. 15070063, 100U/mL;100g/mL) and B-mercapto-ethanol (Sigma-Aldrich, cat. no. M6250, 0.05mM); at 37 °C in 5% CO_2_. Cells were regularly passaged with Trypsin-EDTA (Sigma Aldrich, cat. no. T4174) solution onto pre-coated 6-well plates. Plates were pre-coated with cold high-glucose DMEM (Gibco, cat. no. 41965-062), Penicillin/Streptomycin (Gibco, cat. no. 15070063), Fibronectin (Sigma-Aldrich, cat. no. F1141), and ECM gel (Sigma Aldrich, cat. no. E1270) and incubated at 37 °C for 1h.

### Methods details

#### Data retrieval and integration

We generated an integrated single-cell RNA-seq dataset of 12 published SMART-seq and SMART-seq2 datasets of human pancreatic islets. FASTQ files were downloaded from Wang et al. (GSE83139), Xin et al., (GSE81608), Enge et al. (GSE81547), Segerstolpe et al. (E-MTAB-5061), Lawlor et al. (GSE86469), Li et al. (GSE73727), Camunas-Soler et al. (GSE124742), Dai et al. (GSE164875), Liu et al (GSE270484), Marquez-Curtis et al. (Human Cell Atlas), and SMART-seq and Patch-seq datasets from the Human Pancreas Analysis Program (HPAP)^13–22^. The datasets were obtained via the SRA toolkit (NCBI), ArrayExpress (EMBL-EBI), the Human Cell Atlas, or PANC-DB.

All FASTQ files (one file per donor) were processed simultaneously by removing Illumina sequencing adapters and low-quality bases by TrimGalore (version 0.0.6, default parameters) and quality check were performed using FastQC (version 0.11.9). The reads were mapped to the *Homo sapiens* GRCh38 reference genome (Ensembl GTF version 102) using STAR (version 2.7.7a), which was indexed with sjdOverhang values of 49, 74 or 99 depending on the FASTQ read length of the sample, to produce a BAM sequence alignment file for each donor. Multimappers were filtered out using Samtools (version 1.16.1). The BAM files were directly used for transcriptome analysis and variant calling. In case of paired-end sequenced samples, we used R1 reads for transcriptome analysis, and we used two separate BAM files with either R1 or R2 reads for variant calling.

#### Transcriptome analysis

BAM alignment files were split into one BAM file per cell, and the reads falling in the exon region were counted with FeatureCounts (version 2.0.1). Single-cell gene count tables were filtered based on the following criteria: (1) the minimal number of reads per cell was set to 1×10^4^; (2) cells with mitochondrial gene fractions below 0.25 were kept; (3) the minimum number of genes per cell was set to 200 and applied to all donors. Furthermore, genes that were found in less than three cells were removed from downstream analysis. Single-cell RNA-seq analysis and visualization were done with the Python package Scanpy (version 1.9.3). Doublets were removed using the python package scrublet (version 0.2.3) by filtering out cells with a doublet score above 0.3. Reads were normalized to 1×10^4^ and logp-transformed.

The total counts, the percentage of mitochondrial genes, sjdbOverhang values used in STAR, and the dataset were regressed out using the Scanpy function regress_out() to limit their effect during downstream analysis. The data was batch corrected and dimensionally reduced using principal component analysis (PCA) by harmonypy (version 0.0.10). This was followed by computing of the Euclidian distance between nearest neighboring cells on the Harmony-adjusted PCA space using the neighbors() function in scanpy with n_neighbors set to *√n cells* rounded to the nearest integer, and n_pcs set to 50. Cells were projected in a uniform manifold approximation and projection (UMAP) and clustered using the Leiden algorithm with the resolution set to 1.25. Leiden clusters were annotated by inspecting the expression of conventional pancreatic marker genes.

#### Variant calling and filtering

The Best Practice pipeline from the Genome analysis toolkit (GATK, version 4.1.9.0) was used to call and filter donor-specific SNPs and indels^62^. Pseudo bulk donor-specific BAM files (i.e., all single-cell reads merged into one BAM file) were used as input. Some donors were included in multiple studies; in those cases, we merged the study-specific BAM files into a single donor-BAM file. In short, the duplicated aligned reads were removed, and the sequencing quality score was recalibrated. Variants were called by HaplotypeCaller, and to increase confidence in the discovery of documented rare and common SNPs, the dbSNP VCF (dbSNP Build Number 151, GRCh38.p7) file from NCBI was used as a reference. GVCF files containing unfiltered variants from each donor were merged into a single cohort GVCF file using CombineGVCFs for joint genotyping and variant filtering by GenotypeGVCFs with default filtering parameters. To select for 3′ UTR-specific variants, a custom GTF feature file was made by collapsing multiple gene-specific 3′ UTR entries into a single 3′ UTR entry per gene by selecting the most 5′ stop codon as the start position and using the end position of the farthest 3′ UTR from the Ensembl GTF file (version 102). 3′ UTR variants were selected by intersecting the VCF file with the custom GTF file using bedtools (version 2.30.0). Using the vcf-annotate tool from VCFtools (version 0.1.13), VCF variants were annotated with gene symbols extracted from the GTF file. Rare variants were excluded by applying a minimum population minor allele frequency (MAF) threshold determined by the smallest group in the analysis (T2D, *n* = 35). Assuming Hardy-Weinberg equilibrium, the expected carrier fraction is 2*pq* + *q*^2^ = 1 − (1 − *q*)^2^, where *q* is the MAF and *p* = 1 − *q*. Requiring ≥5 carriers yields a threshold of 0.07. Empirical population MAFs were obtained from the 1000 Genomes Project^63^ (release 20181203, biallelic SNVs) using PyVCF (version 0.6.8), and variants with MAF < 0.07 were excluded from downstream analyses.

#### Gene sets retrieval

Gene sets were retrieved from Gene Set Enrichment Analysis (GSEA) database^64^. We used the beta identity gene sets from van Gurp et al.^65^ and the Hallmark unfolded protein response (v2026). RNA-binding protein (RBP) gene symbols were retrieved from the CISBP-RNA database from the MEME Suite^36^ and Van Nostrand et al.^37^. All genes included per gene set are reported in Table S3.

#### RNA isolation

Frozen islets (Table S4) were received through the Integrated Islet Distribution Program (IIDP). The AllPrep DNA/RNA/miRNA Universal kit (Qiagen, cat. no. 80224) was used to isolate the total RNA of cell lysates of isolated human islets and EndoC-βH1 cells according to the manufacturer’s protocol. After, RNA was measured for purity and concentration on the Nanodrop. Total RNA was subsequently used for small RNA-seq, and miRNA RT-qPCR.

#### Small RNA-seq data analysis

A sequencing library was generated from total isolated RNA from isolated human islets using the D-Plex Small RNA-seq Kit (Diagenode, cat. no. C05030001) for paired-end Illumina sequencing (100 cycles).

Only R1 FASTQ files were used for downstream analysis. To process the FASTQ files for analysis, first, UMI was extracted and appended to the end of its corresponding read name using fumi_tools (version 0.12.2) copy_umi, and reads were trimmed for UMIs, sequencing adapters, and poly(A) tail by cutadapt (version 3.4). Only trimmed reads between 15 and 30 nucleotides were retained. The reference genome was indexed using STAR with sjdbOverhang set to 1, incorporating the GENCODE annotation (release 24). Reads were aligned to the *Homo sapiens* GRCh38 reference genome using STAR with a custom miRNA-specific annotation GTF file provided by Hita et al^66^, and following parameters from the ENCODE miRNA-seq workflow. End-to-end alignment with stringent mismatch filtering was used, with splice junction detection disabled to accommodate short miRNA reads. Limited multimapping was allowed, and transcriptome-level outputs were generated. Transcriptome-aligned BAM file reads were deduplicated using fumi_tools. miRNA-level read counts were generated using a custom Python script. Reads mapping to multiple miRNAs were counted as fraction 1/N where N is the number of mapped miRNAs. Islet samples with less than 2.75×10^5^ raw counts were considered outliers and were removed from the analysis. Samples passing this QC are reported in Table S4. miRNAs detected in fewer than two samples or with fewer than five raw counts were removed. Counts were size-factor normalized using the DEseq method by the R package DeSeq2 (version 1.38.3). Principal Component Analysis (PCA) was performed using the sklearn.decomposition.PCA python package using variance-stabilized counts. Heatmap plots were generated using PyComplexHeatmap (version 1.7.7).

#### miRNA and RBP binding prediction

22 nucleotide sequences upstream and downstream flanking the eQTLs were extracted in FASTA format for both the reference and alternative alleles from the *Homo sapiens* GRCh38 reference genome (Ensembl GTF release 102). This extraction was performed using a combination of bedtools, bcftools, and custom Python scripts. The FASTA file was used for miRNA and RBP binding prediction. For miRNA binding prediction, mature miRNA sequences in FASTA format were downloaded from miRbase (release 22.1)^67^. miRNA binding scores were computed using the miRanda software^48^ using the strict seed binding-mode, with the miRNA FASTA and eQTL FASTAs as input. The settings applied were: score threshold of 50, energy threshold of 1 kcal/mol, scaling parameter of 4, gap-open penalty of -4, and gap-extend penalty of -9. miRNA-eQTL pairs were selected if only one of the alleles led to strict seed binding. Only miRNA-eQTL pairs where the binding allele led to a significant reduction of transcription, as defined by the eQTL effect size direction, compared to the other alleles were kept. Heatmap plots were generated using PyComplexHeatmap (version 1.7.7).

For RBPs, MEME Motif Format files were either directly downloaded from the CISBP-RNA database from the MEME Suite or converted from logo images retrieved from van Nostrand et al. 2020^37^ using Logo2PWM^68^ in MATLAB (version R2023b). Binding prediction was performed using FIMO (The MEME Suite) with the p-value threshold set to 1e-2 and the MEME Motif Format files, eQTL FASTAs, and a custom background model file as input. This background file was built using a FASTA file containing all human 3′ UTRs using fasta-get-markov. The input FASTA file for this background was generated with bedtools getfasta with the -s option enabled. RBP binding gain/loss was considered when one of the alleles leads to an alteration or loss of binding.

#### miRNA RT-qPCR

Isolated total RNA from EndoC-βH1 cell lysates was used as a template for reverse transcription. TaqMan™ MicroRNA Reverse Transcription Kit (Applied Biosystems, cat. no. 4366596) was used to perform reverse transcription according to the manufacturer’s instructions, with the assays/probes: RNU48 (Applied Biosystems, cat. no. ID001006), and hsa-miR-95 (Applied Biosystems, cat. no. ID000433). miRNA abundance was quantified with the TaqMan™ Fast Advanced Master Mix for qPCR (Applied Biosystems, cat. no. 4444556) and the above-mentioned probes on the Bio-Rad CFX384 Real-Time PCR system. miR-95-3p Cycle threshold (Ct) values were normalized to RNU48 Ct values.

#### miRNA transfections

Approximately 1-1.5×10^5^ EndoC-βH1 cells were seeded onto 48-well plates (Greiner, cat no. 677180) and cultured overnight. Lipofectamine RNAiMAX (Invitrogen, cat. no. 13778100) was used for the transfection of cells with varying concentrations (10 nM, 20 nM, 30 nM) of hsa-miR-95-3p mirVana® miRNA inhibitor (Invitrogen, cat. no. 4464084) and mirVana® miRNA mimic (Invitrogen, cat. no. 4464066). As negative control we used 30 nM of mirVana™ miRNA Mimic Negative Control #1 (Invitrogen, cat. no. 4464058). Following the transfections, cells were cultured for 48h.

#### *QDPR* RT-qPCR

RNA was isolated using Trizol extraction and used as a template for reverse transcription, which was performed with Superscript III reverse transcriptase (Invitrogen, cat. no. 18080093) and oligo(dT) primers (Integrated DNA Technologies). *QDPR* transcript abundance was quantified with iQ SYBR Green Supermix (Bio-Rad, cat. no. 1706686) and *QDPR* forward primer (5′-GTGACTGCTGAGGTTGGAAAGC-3′) / *QDPR* reverse primer (5′-CTCTGCTTCCACATCAGGTCAC-3′) combination. Ct values were normalized with *GAPDH* transcript abundance, measured with *GAPDH* forward primer (5′-GGAGCGAGATCCCTCCAAAAT-3′) and *GAPDH* reverse primer (5′-GGCTGTTGTCATACTTCTCATGG-3′) primers. All the measurements were performed on the Bio-Rad CFX384 Real-Time PCR system. *GAPDH* normalized Ct values were normalized to the non-transfection control.

#### Sanger sequencing genotyping

EndoC-βH1 cells were genotyped for rs1049581/rs1049582 (*QDPR*) from genomic DNA (gDNA), isolated using the DNeasy Blood & Tissue Kit (Qiagen, cat. no. 69504). Isolated gDNA was checked for purity and concentration on Nanodrop. The *QDPR* 3′ UTR region was PCR amplified from gDNA using Q5 polymerase (New England Biolabs, cat. no. M0491L) using the *QDPR* forward primer (5′-CTTTTAAACAGTCGCTGCTGTG-3′) and reverse primer (5′-GCAACCACACAGGAAAAGGATA-3′) combination. The PCR amplicon was isolated with the QIAquick PCR Purification Kit (Qiagen, cat. no. 28104) according to the manufacturer’s protocol and measured for purity and concentration on the Nanodrop. Sanger sequencing was performed with a *QDPR* (5′-AAGGAAGGACGGAACTCACCCC-3′) sequencing primer by Macrogen.

#### HumanIslets.com dataset

We used measurements, including bulk mRNA-seq (European Genome-phenome Archive: EGAS00001007241), static, and dynamic GSIS on primary human islets from the HumanIslet.com project that were used in the study of Kolic et al^35^. Donors were genotyped by running a custom script that extracts 3′ UTR eQTL genotypes directly from bulk mRNA-seq BAM files using samtools. Variants with a minimal read depth of 10 were genotyped using a binomial-likelihood model in Python, comparing the observed alternative allele counts against homozygous reference, heterozygous, and homozygous alternative states, assuming a sequencing error rate of ε = 0.01 and assigning the genotype with the highest log-likelihood. Regression analysis between allele dosage and GSIS values was performed using the R package limma (version 3.54.2). For dynamic GSIS, raw secretion values were normalized to pmol/L/islet after subtraction of the 3 mM glucose baseline. Insulin secretion was then separated by stimulation phase. For the first phase, peak secretion was defined as the maximum observed value. For both the first and second phases, the area under the curve (AUC) was calculated using the trapz function from the NumPy Python package (version 1.23.5). Donor age, sex, BMI, and Hb1Ac, islet purity, islet alpha cell content, and islet beta cell content were included as covariates. Extreme outliers were removed by log10-transforming positive values and excluding data points exceeding four standard deviations above the mean on the log scale. P-values were adjusted using the Benjamini-Hochberg False Discovery Rate (FDR) procedure (Python statsmodels package, version 0.14.1). Adjusted p-values below 0.05 were considered significant.

#### Sorted beta cell mRNA-seq

RNA-seq data from sorted beta cells were downloaded from PANC-DB^50^ and aligned to the *Homo sapiens* reference genome (GRCh38) using STAR (version 2.7.11b). Gene-level read counts were quantified with featureCounts (version 2.1.1) based on Ensembl gene annotations (v115). Counts were normalized to transcripts per million (TPM) to account for differences in gene length and sequencing depth, enabling quantitative comparison of gene expression across samples. Donors diagnosed with any form of diabetes and ND donors present in the scRNA-seq dataset were excluded. Genes scores were calculated per gene set (Table S3) in R using the simpleScore function from Singscore (version 1.18.0) and AddModuleScore from Seurat (version 5.4.0). Regression analysis was performed using limma (version 3.54.2) with donor age, sex, and BMI, included as covariates.

## Quantification and statistical analysis

Conditions were compared using an independent Student’s t-test (SciPy.stats), and p-values were adjusted using the Bonferroni method (statsmodels), unless specified. Fisher’s exact test (SciPy.stats) was used for enrichment analyses. P-values below 0.05 were considered statistically significant and are shown in the figure panels or in the text. Furthermore, all eQTL statistical analyses are presented in Table S2. For experimental assays, data are presented as ±SEM, and the number of biological replicates is mentioned in the figure legends. In some figure panels, other statistical methods were used, which are described below.

### Cis-eQTL computation

Cis-eQTLs were computed per cell type and T2D status using zero-inflated negative binomial regression, adapting the approach implemented in the R package SCeQTL (version 0.2.0)^24^. For each variant-gene pair, DESeq2-normalized gene expression was modeled as a function of genotype dosage (0/1/2) while adjusting for donor- and cell-level covariates. The count component included genotype dosage and the covariates age, sex, study, log(total counts), percent mitochondrial reads, and the first five gene-level principal components. The zero-inflation component used the same covariates.

Full model:

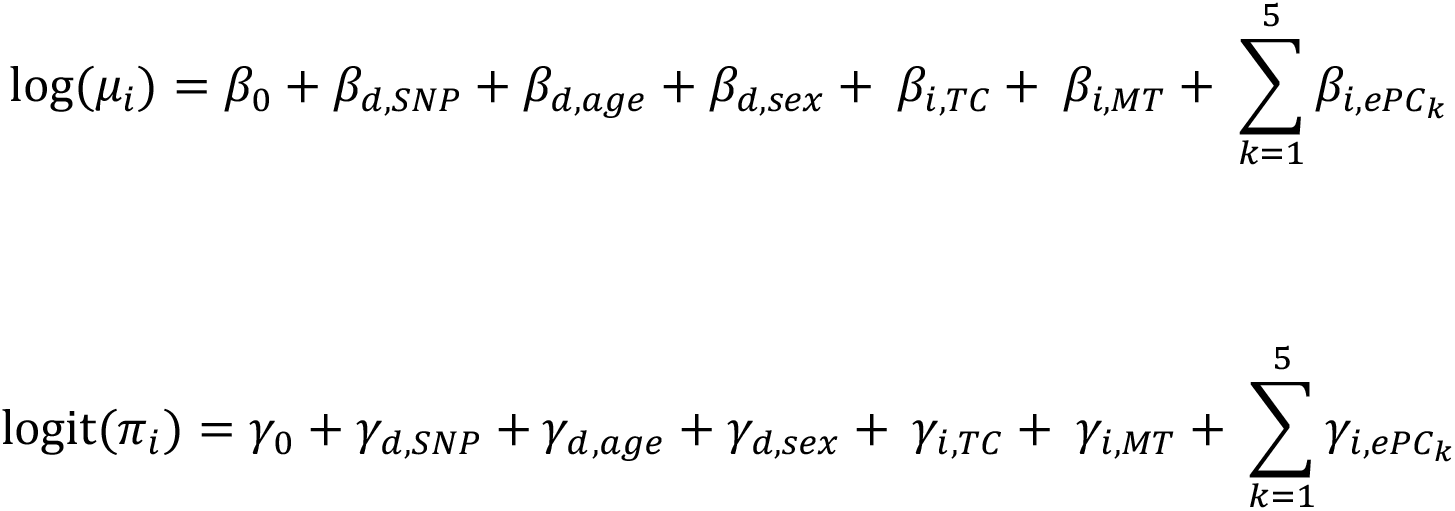

nested model:

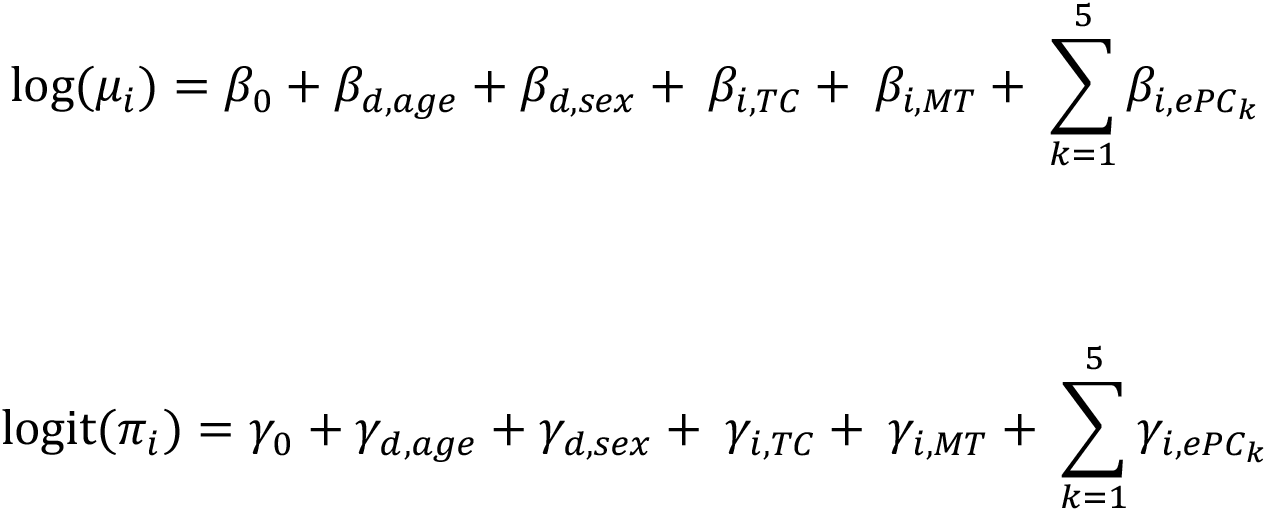

Models were fitted using zeroinfl from the R package pscl (version 1.5.9), assuming a negative binomial distribution, with EM estimation enabled and a maximum of 300 iterations. As in SCeQTL, the significance of the genotype effect was assessed by a likelihood-ratio test comparing the full model (including genotype) to a nested null model (excluding genotype), with p-values obtained from a chi-squared distribution with 1 degree of freedom. Tests were restricted to variant-gene pairs with at least 10 cells with non-missing genotype dosage and at least two genotype classes present. To reduce the influence of extreme non-zero expression values, outlier filtering was applied by removing observations exceeding the median of non-zero expression plus four median absolute deviations (MAD). P-values were adjusted using the FDR procedure (Python statsmodels). To account for noise due to lowly expressed genes, we excluded genes with a cell type-specific median normalized expression of 0.4 or expression in less than 20% of cells. We further restricted eQTL effects to those exhibiting a monotonic directionally consistent dosage-expression relationship, defined as strictly increasing or decreasing genotype-stratified expression medians with a matching sign of the genotype effect (β). Associations with FDR-adjusted p-values < 0.05 were considered significant, and β was used to assess allelic directionality. Due to the relatively low number of donors, we focused only on general eQTL effects and did not investigate sex differences.

The Fisher’s exact test (SciPy.stats version 1.10.1) was used to assess differences in eQTL counts between T2D states and was restricted to variants passing QC in both groups and mapping to shared genes. eQTL overlap with bulk tissue datasets was assessed by intersecting genomic positions with eQTLs from all tissues in the GTEx v8 database^28^ and from whole islets in the TIGER database^27^. For GTEx, tissues belonging to the same organ system were grouped, and the percentage of overlapping eQTLs was averaged across tissues within each group, with the standard deviation reported.

### Linkage disequilibrium analysis

Biallelic SNVs from the 1000 Genomes Project^63^ (release 20181203) were used to calculate linkage disequilibrium (LD) between 3′ UTR eQTL variants and variants located in promoter regions or active islet enhancers. LD was computed with PLINK (version v2.0.0-a.7LM) using the r2-unphased function within a 100 kb window. Variants showing high LD (*r*^2^ > 0.8) with each 3′ UTR eQTL variant were intersected with promoter and enhancer annotations using bedtools. Promoter regions were defined from transcription start site annotations provided by the Eukaryotic Promoter Database^69^, whereas active islet enhancers were derived from imputed epigenomic annotations from the NIH Roadmap Epigenomics Mapping Consortium^70^.

### SNP eQTL GWAS-colocalization

GWAS summary tables for T2D risk were retrieved via the DIAGRAM consortium^29^, and glycemic traits (i.e., fasting glucose, fasting insulin, glucose 2 hours post oral glucose tolerance test, and HbA1c) were retrieved via the MAGIC consortium^30^. Colocalization was performed per cell group using the R package coloc (version 5.2.3) using the coloc.abf function with the eQTL and GWAS effect sizes. The analysis was performed per 3′ UTR, and we included all variants regardless of MAF or monotonicity. Colocalization was considered for PP.H4 > 0.8.

### Differential gene and gene set analysis

Differential gene expression analysis was performed with the Wilcoxon Signed Rank Test using the rank_genes_groups function in Scanpy. We considered genes with an adjusted p-value below 0.05 as differentially expressed. Gene set enrichment analysis was performed with GSEApy (version 0.10.8), including the GO Biological Process (v2025) and MSigDB Hallmark (v2020) databases.

### Motif enrichment analysis

BED files containing annotated AU, GU, U-rich elements (ARE, GRE, URE), and polyadenylation signals (PAS) motifs were retrieved from the AREsite2 database^41^. For each motif set, strand-aware distances from 3′ UTR variants to the nearest motif within the same 3′ UTR were computed using bedtools closest with the options -s and -D ref. Motifs within a window of ±50 nucleotides up- and downstream of variants were retained. Motif enrichment was assessed by comparing the observed eQTL distance probability distribution to a null distribution generated by permutation: for each motif, the same number of background variants (non-eQTLs) as eQTLs was randomly sampled 5,000 times and binned into 5-nucleotide distance bins. An empirical enrichment plus-one corrected p-value was computed for each positional bin as the fraction of permutations with background density ≥ observed eQTL density using p = 0.05 as the significance threshold.

### Differential miRNA analysis

Differential miRNA expression was computed using the DESeq2 (version 1.38.3) package in R. For each differentially expressed miRNA, Pearson correlation was performed between miRNA expression levels and BMI or HbA1c, and p-values were adjusted using the FDR procedure (Python statsmodels).

## Additional resources

### Data availability

Public human pancreatic islet Smart-seq and Smart-seq2 datasets can be retrieved from the NCBI Gene Expression Omnibus (GEO) under the accession numbers GSE83139, GSE81547, GSE81608, GSE86469, GSE73727, GSE124742, GSE164875, GSE270484, the EMBL-EBI ArrayExpress database under accession number E-MTAB-5061, the Human Cell Atlas, and PANC-DB (https://hpap.pmacs.upenn.edu). Glucose-stimulated insulin secretion data is available through humanislets.com. Islet bulk mRNA-seq is available through European Genome-phenome Archive (EGA) under the accession number EGAS00001007241. Sorted beta cell RNA-seq is available through PANC-DB (https://hpap.pmacs.upenn.edu). Small RNA-seq data is available through EGA under accession number EGAD50000002342.

### Code availability

Scripts and notebooks for the eQTL pipeline are available at https://github.com/tjj-de-winter/3UTReQTL_islets, and scripts for smallRNA-seq processing are available at https://github.com/tjj-de-winter/miRNA_mapping

### Ethical statement

This work includes data and/or analyses from HumanIslets.com funded by the Canadian Institutes of Health Research, JDRF Canada, and Diabetes Canada (5-SRA-2021-1149-S-B/TG 179092) with data from islets isolated by the Alberta Diabetes Institute IsletCore with the support of the Human Organ Procurement and Exchange program, Trillium Gift of Life Network, BC Transplant, Quebec Transplant, and other Canadian organ procurement organizations with written informed donor consent as approved by the Human Research Ethics Board at the University of Alberta (Pro00013094).

## Funding

Financial support for this study was mainly provided by the DON Foundation, Dutch Diabetes Research Foundation, and Nederlandse Organisatie voor Wetenschappelijk Onderzoek (NWO, OCENW.XL.23.084). In addition, EdK and FC have received funding from the Bontius Foundation, Tjanka Foundation, Novo Nordisk Foundation Center for Stem Cell Medicine reNEW (NNF21CC0073729) and RegMedXB. AA is supported by the European Research Council Starting grant (ERC-StG 101042634-i-Signaltrace), the Nederlandse Organisatie voor Wetenschappelijk Onderzoek (NWO) VIDI grant (VI.Vidi.203.050), and La Caixa Foundation (CaixaResearch Health 2023 Call #COMET-HR23-00392). Work at Stanford was supported by NIDDK (UM1-1DK126185; U01-DK123716; U24-DK138512) and the Wellcome Trust (200837).

## Acknowledgements

We extend our sincere gratitude to the pancreas donors and their families. We also thank Diagenode, and the Looijenga group of the Prinses Máxima Centrum for their contributions. This manuscript used data acquired from the database (https://hpap.pmacs.upenn.edu/) of the Human Pancreas Analysis Program (HPAP; RRID:SCR_016202; PMID: 31127054; PMID: 36206763; PMID: 35228745; PMID: 37188822). HPAP is part of a Human Islet Research Network (RRID:SCR_014393) consortium (UC4-DK112217, U01-DK123594, UC4-DK112232, and U01-DK123716). Lastly, we thank the lab members from the Alemany, de Koning, and Geijsen groups for their valuable input and discussions.

## Author contributions

TdW and AA designed the study, with input from FC and EdK. TdW integrated and analyzed the scRNAseq and the smallRNAseq data; developed the eQTL detection pipeline; and analyzed all experiments, with input from AA, FC, and EdK. MS performed the cell line experiments. HS performed the bioinformatic analysis on the data from the HumanIslets.com project and sorted beta cells, under the supervision of PM, JJ, and ALG. PM and ALG contributed to the acquisition of public scRNAseq data. TdW and AA wrote the manuscript, with input from ALG, FC, and EdK. All authors read and approved the manuscript.

## Declaration of interest

ALG’s spouse is an employee of Genentech and holds stock options in Roche.

## Declaration of generative AI and AI-assisted technologies in the writing process

During the preparation of this work, the authors used ChatGPT to improve the readability, clarity, and language. After using this tool/service, the authors reviewed and edited the content as needed and take full responsibility for the content of the published article.

## Supplementary table captions

**Supplementary Figure S1.**
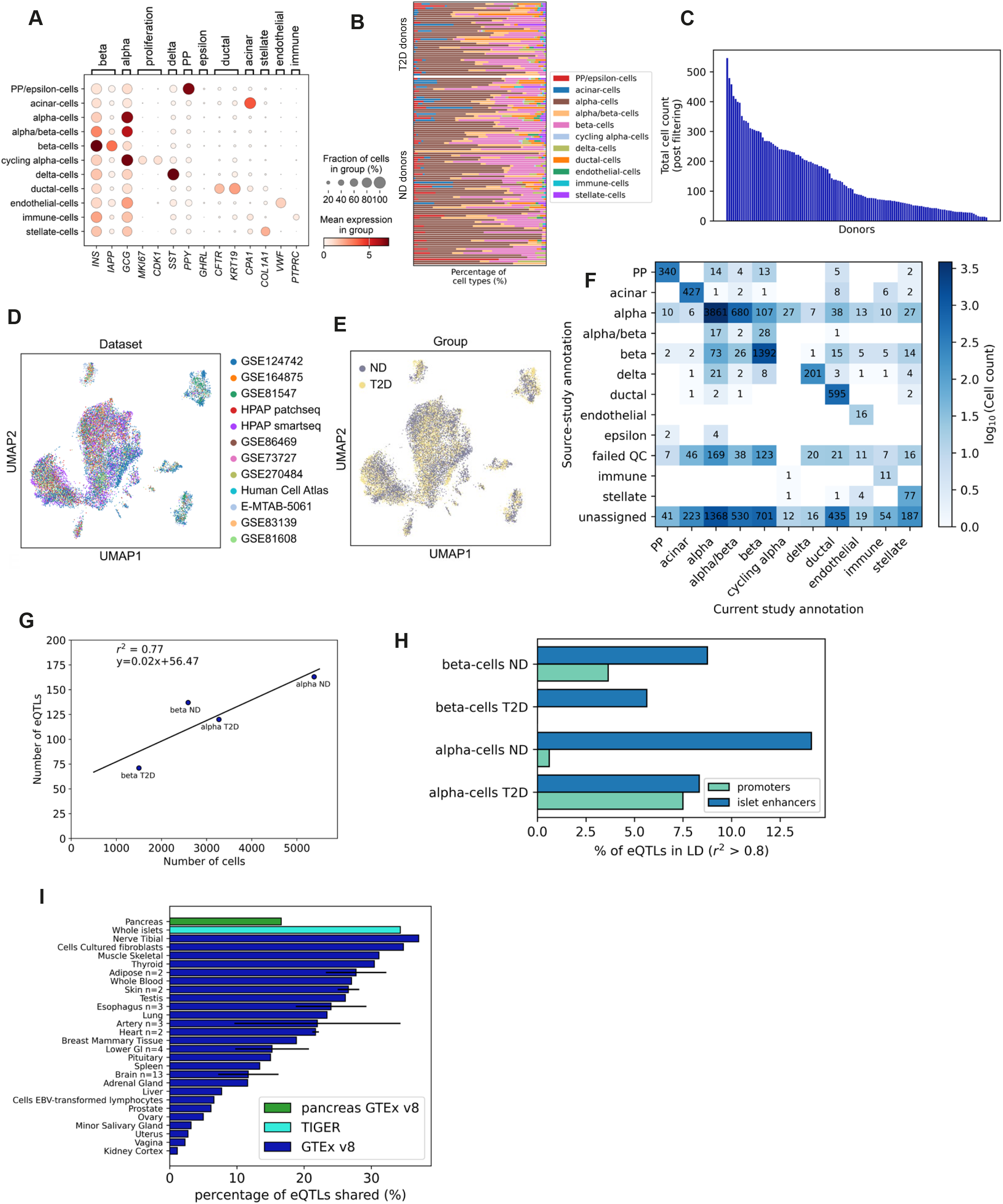
**A**, Dotplot showing the expression of canonical cell type markers for each cell type cluster. **B**, Stacked bar graph containing the fraction of cell types per donor for the ND and T2D cohort. **C,** Bar graph showing the number of cells per donor post filtering. **D**,**E**, UMAP where cells are colored according to the dataset origin (D) or diabetes status (ND or T2D) (E). **F**, Heatmap showing concordance between cell-type annotations from the original source-study and the annotations assigned after integration and reanalysis in the current study. Only cells with available source-study annotations are shown. Numbers indicate cell counts for each annotation pair, with color intensity representing the log_10_-transformed cell count. **G**, Scatter plot showing the correlation between the number of cells per cell type and the number of detected eQTLs. The solid line indicates the result of a linear fit to the displayed data, and *r*^2^ value measures the goodness of the fit. **H**, Bar plot showing the percentage of cell type-specific 3′ UTR eQTLs in high LD (*r*^2^ > 0.8) with promoter and islet enhancer variants **I**, Bar plot showing the percentage of cell type-specific eQTLs also present in other tissues (GTEx and TIGER databases). Tissues with error bars indicate organs with multiple tissues merged, and these bars represent mean overlap ± SEM, with the number of tissues included in the y-label.

**Supplementary Figure S2.**
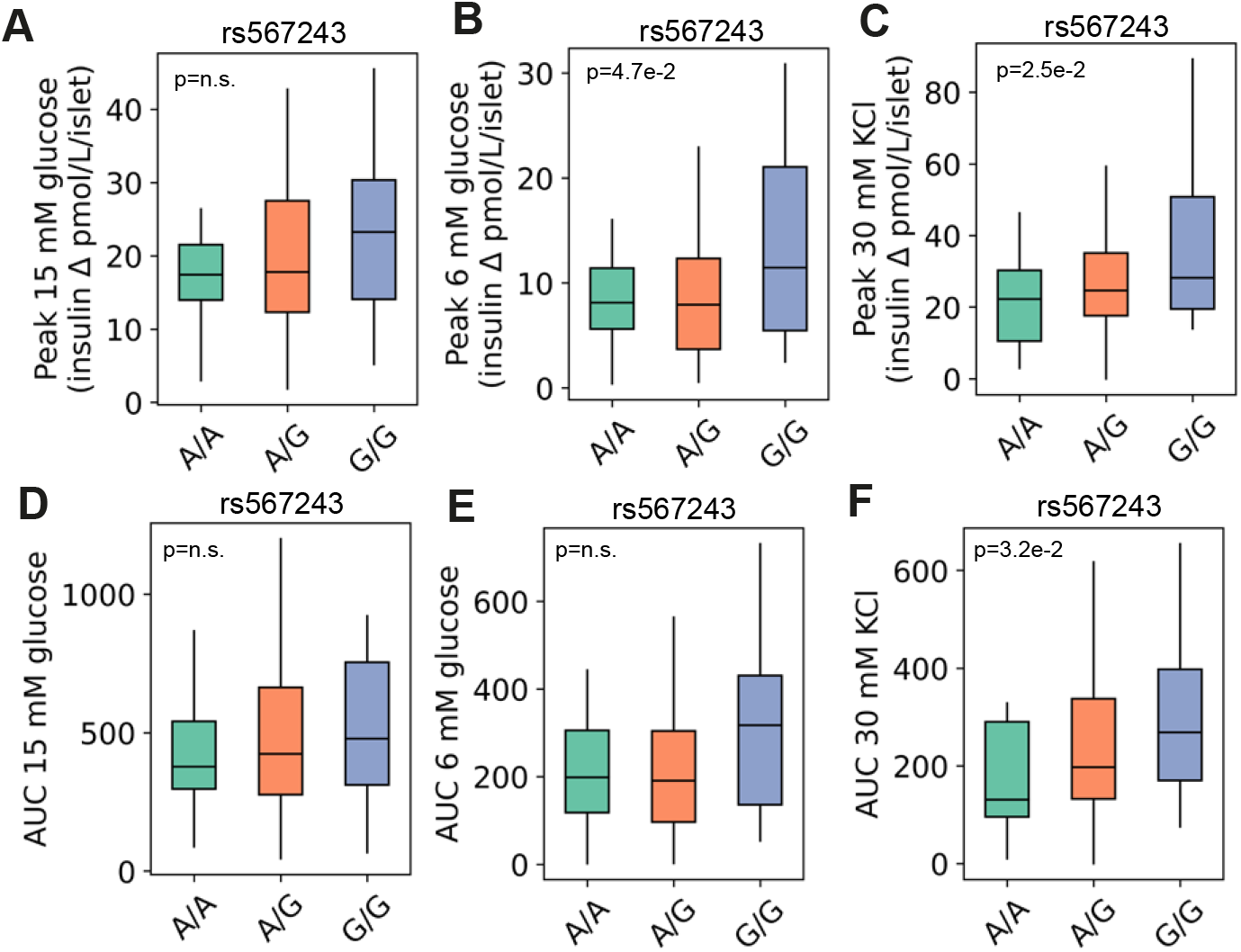
**A**-**C**, Genotype-stratified boxplot showing the peak insulin secretion (first-phase) per stimulus of ND donor islets assessed using dynamic GSIS (basal glucose, 3mM; high glucose, 15 mM; moderate glucose, 6 mM; 30 mM KCl) by genotype of *G6PC2* variant rs567243. The plot is annotated with the p-value from linear regression adjusted for donor and islet isolation-level covariates. Center line: median, box: interquartile range, vertical lines: spread of the data. 15 mM glucose stimulus (A), 6 mM glucose stimulus (B), 30 mM KCl stimulus (C). **D**-**F**, Genotype-stratified boxplot showing the area under the curve (AUC) (first- and second-phase) per stimulus of ND donor islets assessed using dynamic GSIS (basal glucose, 3mM; high glucose, 15 mM; moderate glucose, 6 mM; 30 mM KCl) by genotype of *G6PC2* variant rs567243. The plot is annotated with the p-value from linear regression adjusted for donor and islet isolation-level covariates. Center line: median, box: interquartile range, vertical lines: spread of the data. 15 mM glucose stimulus (D), 6 mM glucose stimulus (E), 30 mM KCl stimulus (F).

**Supplementary Figure S3.**
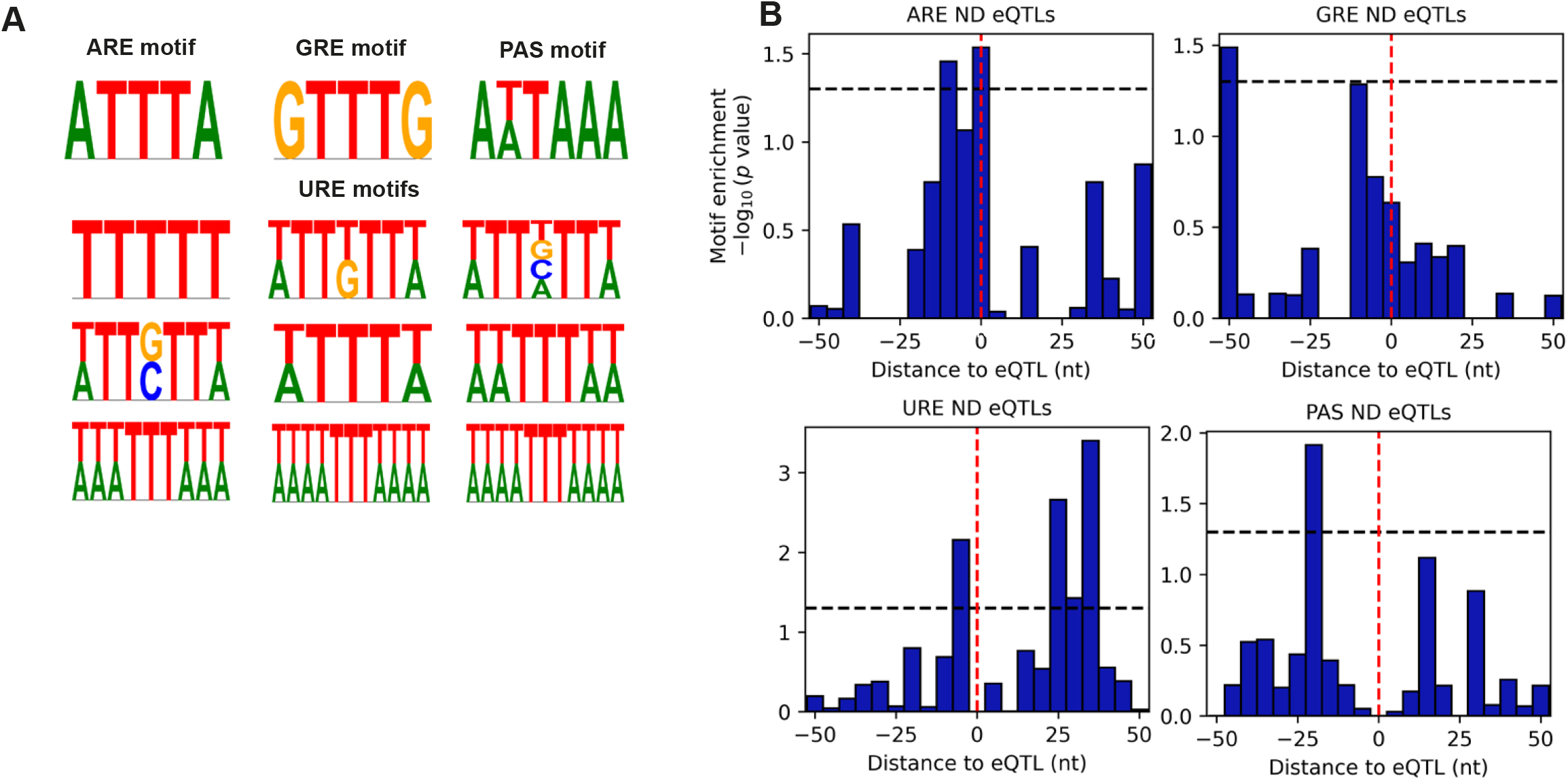
**A**, DNA sequence logo plots showing motifs of 3′ UTR regulatory elements: AU-rich element (ARE), U-rich element (URE), G-rich element (GRE), and polyadenylation signal (PAS). **B**, Bar graphs showing motif enrichment (−log_10_ p-value) for ARE, GRE, URE, and PAS, within a 50-nt window around eQTL variants in ND donors. The window is binned in 5-nt, 0-centered intervals (red dashed line indicates eQTL position). The horizontal dashed line marks significance (p < 0.05).

**Supplementary Figure S4.**
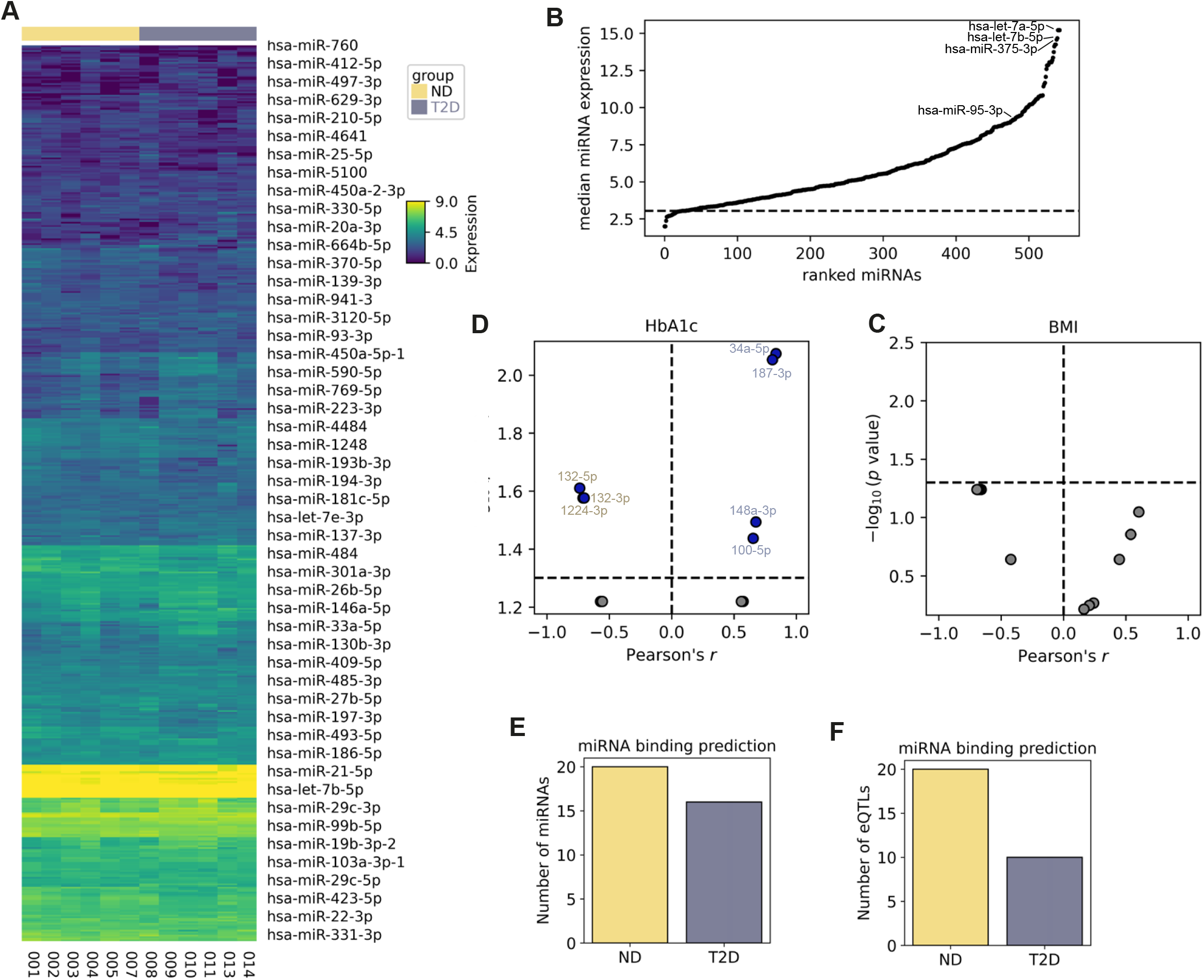
**A**, Heatmap showing log_2_(normalized expression) levels of all detected miRNAs in human pancreatic islets per donor. Donors are annotated by their T2D status. **B**, Rank plot showing median miRNA expression across donors for all detected miRNAs. The dashed line indicates the 5th percentile cutoff, and miRNAs above the line represent the top 95% by median expression. The top miRNAs and miR-95-3p are annotated. **D**,**C**, Correlation volcano plots of associations between miRNA expression and BMI (D) and HbA1c (C) for differentially expressed miRNAs, showing Pearson’s r versus −log_10_(BH-adjusted p-value). The vertical dashed line marks no correlation (r=0). The horizontal dashed line marks significance (p < 0.05). Significant miRNAs are annotated, and the annotation color (yellow = ND; gray = T2D) indicates the cohort in which the miRNA is upregulated. **E**,**F**, Bar graphs showing the number of miRNAs (E) and eQTLs (F) for predicted miRNA-eQTL pairs with disrupted binding sites.

**Supplementary Figure S5.**
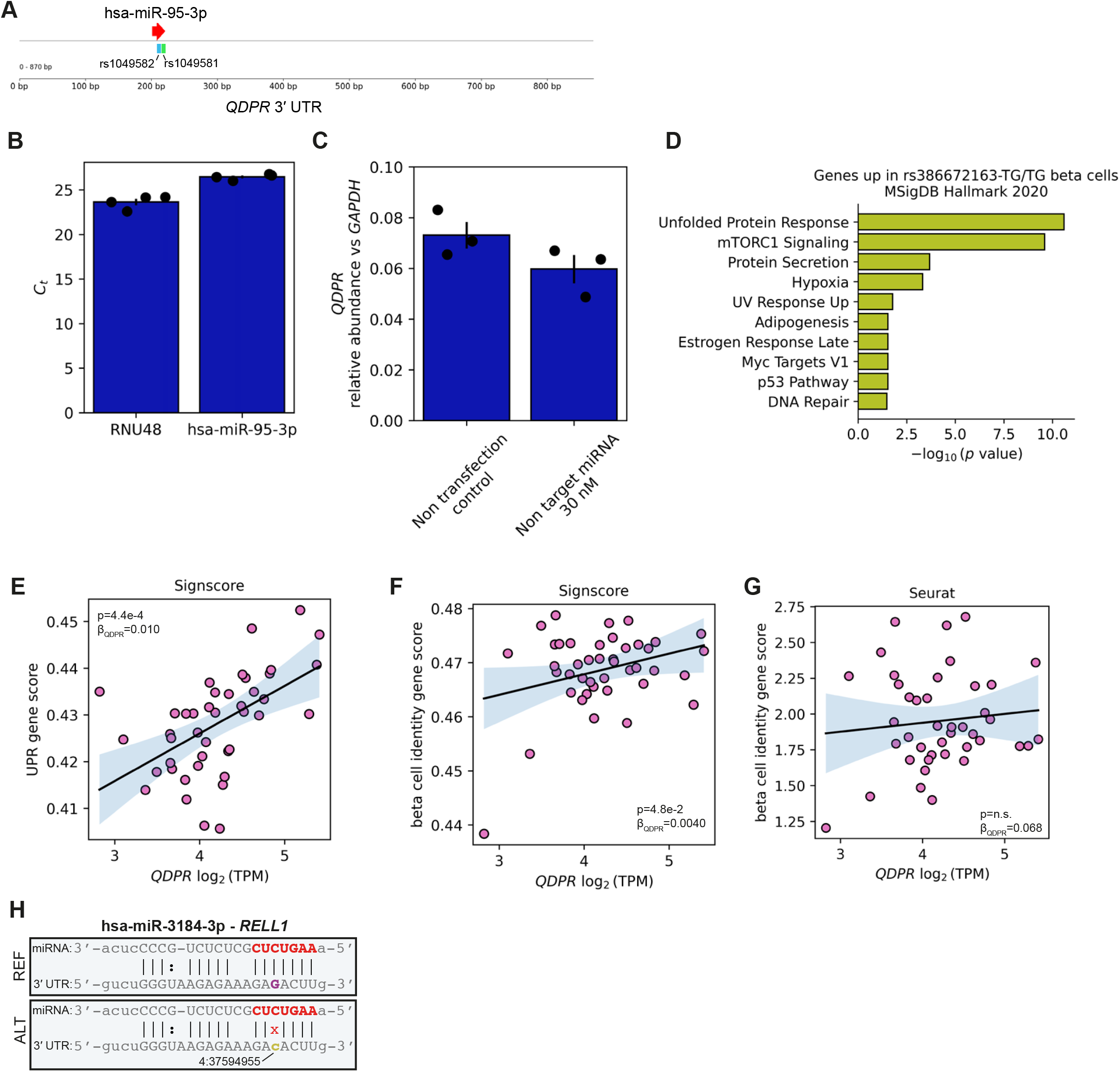
**A**, Overview of the *QDPR* 3′ UTR showing a single predicted binding site of miR-95-3p overlapping the variants rs1049581 and rs1049582. **B**, Raw Ct values obtained from miRNA RT-qPCR for the housekeeping snoRNA RNU48 and miR-95-3p in EndoC-β1. N = 4 biological replicates, mean ± SEM. **C**, *GAPDH* normalized *QDPR* abundance in EndoC-β1 measured by RT-qPCR for the non-transfection and non-targeting miRNA controls. N = 3 biological replicates, mean ± SEM. Each point represents the average of three wells. **D**, Top 10 results from gene set enrichment analysis (MSigDB Hallmark) using upregulated genes in beta cells with the alternative rs386672163 TG/TG genotype from Figure 5G. **E**-**G** Scatter plots showing the association between log_2_(normalized *QDPR*) expression versus UPR (E) and beta cell identity gene scores (F, G). The gene scores were calculated using Signscore (E,F) or Seurat (G) in sorted beta cells, and plots are annotated with the effect size (β) and p-value obtained from regression analysis adjusted for donor-level covariates. The solid line indicates the result of a linear fit to the displayed data. **H**, Nucleotide alignment of hsa-miR-3184-3p to the 3′ UTR of *RELL1* shown in the orientation of transcription after Watson and Crick and Wobble base pairing. miRNA seed region shown in bold red, the reference allele (binding) in purple, and the alternative allele (disrupting) in gold, and annotated with the genomic position.

**Supplementary Table S1.** Donor data of islet samples used for eQTL detection, including donor ID, study accession IDs, donor age, sex, BMI, HbA1c, diabetes status, and number of cells in the analysis. This table is related to all figures.

**Supplementary Table S2.** Detected 3′ UTR eQTLs per cell type, including variant genomic position, dbSNP rsID, gene name, reference allele, cell type, donor group, Benjamini-Hochberg (BH) adjusted p-values, and effect size of a zero-inflated negative binomial (ZINB) regression. This table is related to all figures.

**Supplementary Table S3.** Genes included in the RBP, UPR, and beta cell identity gene sets. This table is related to Figures 3, 5, and S5.

**Supplementary Table S4.** Donor data of islet samples used for small non-coding RNA sequencing, including donor ID, age, sex, BMI, HbA1c, isolation center, diabetes history, and which experiments the islets were used for. This table is related to Figures 4 and S4.

**Supplementary Table S5.** Beta cell 3′ UTR eQTLs predicted to have altered miRNA binding sites as computed by the miRanda algorithm, including target miRNA, variant genomic position, gene name, cell type, diabetes status, target allele and locus, miRanda score, binding energy, query binding position, subject binding position, alignment length, subject identity, query identity, eQTL effect size and direction. This table is related to Figures 4 and S4.

**Supplementary Table S6.** Gene set enrichment results based on genes from genotype-aware DGE analysis, including the gene set database, enriched biological process, gene overlap, nominal and adjusted p-values, odds ratio, combined enrichment score and contributing genes. This table is related to Figure 5 and S5.

## Notes

### Summary of Updates

To ensure the robustness of our results, we increased the number of donors from 30 to 125, improved our statistical modelling to account for donor metadata and applied stricter thresholds to identify candidate genetic variants and subsequent 3'UTR eQTLs. In this new version, we propose a computational pipeline that exploits genetic variation in an integrated scRNA-seq dataset of 125 donors to unravel the post-transcriptional regulatory roles of miRNAs and RBPs.

